# Estimating admixture pedigrees of recent hybrids without a contiguous reference genome

**DOI:** 10.1101/2022.12.15.520578

**Authors:** Genís Garcia-Erill, Kristian Hanghøj, Rasmus Heller, Carsten Wiuf, Anders Albrechtsen

## Abstract

The genome of recently admixed individuals or hybrids have characteristic genetic patterns that can be used to learn about their recent admixture history. One of these are patterns of interancestry heterozygosity, which can be inferred from SNP data from either called genotypes or genotype likelihoods, without the need for information on genomic location. This makes them applicable to a wide range of data that are often used in evolutionary and conservation genomic studies, such as low-depth sequencing mapped to scaffolds and reduced representation sequencing. Here we implement maximum likelihood estimation of interancestry heterozygosity patterns using two complementary models. We furthermore develop `apoh` (Admixture Pedigrees Of Hybrids), a software that uses estimates of paired ancestry proportions to detect recently admixed individuals or hybrids, and to find the most compatible recent admixture pedigree. It furthermore calculates several hybrid indices that make it easier to identify and rank possible admixture pedigrees that could give rise to the estimated patterns. We implemented `apoh` both as a command line tool and as a Graphical User Interface that allows the user to automatically and interactively explore, rank and visualize compatible recent admixture pedigrees, and calculate the different summary indices. We validate the performance of the method using admixed family trios from the 1000 Genomes Project. In addition, we show its applicability on identifying recent hybrids from RAD-seq data of Grant’s gazelle (*Nanger granti* and *Nanger petersii*) and whole genome low depth data of waterbuck (*Kobus ellipsiprymnus*) which shows complex admixture of up to four populations.

## 1 Introduction

Hybridization and admixture is a common outcome when different populations or species come into secondary contact, with an increasingly recognized role in evolution, for example during speciation (Feder et al., 2012) or adaptation (Edelman & Mallet, 2021). An important step towards understanding the role of hybridization in natural populations is the ability to reliably detect and characterize the presence of recent hybrids or admixed individuals from genetic data. Hybrid inference based on hybrid indexes or admixture proportions, which model the global proportion of the individual’s genome originating from each admixing population, has a long history in population genetics (Pritchard et al., 2000; Vaha & Primmer, 2006; Buerkle, 2005; Beugin et al., 2018). However, it has been shown that these methods can lead to spurious conclusions regarding admixture (Lawson et al., 2018), and furthermore they do not directly distinguish between very recent hybridization, happening within up to five generations, or admixed individuals where hybridization happened several generations ago. Other methods have harnessed the characteristic signals that admixture leaves in the genomes in the first few generations following the admixture event. One such characteristic pattern of recent hybrids is an excess of regions in the genome where the two alleles at each loci derives from different ancestries. This pattern is known as ‘interancestry heterozygosity’ and is used by multiple existing methods to detect and characterize recent hybrids (Boecklen & Howard, 1997; Anderson & Thompson, 2002; Fitzpatrick, 2012; Crouch & Weale, 2012; Pfaffelhuber et al., 2022; Shastry et al., 2021). Another useful signal to study recent hybridization is the length of continuous stretches of the genome with the same ancestry, known as ‘admixture tracts’. Admixture tract lengths decrease each generation after the admixture event through recombination, and this is used by several methods (Zou et al., 2015; Pei et al., 2020) to learn about recent admixture history. Finally, a combination of both interancestry heterozygosity and admixture tract lengths has been shown to be most powerful, allowing inference of admixture pedigree and time since admixture (Avadhanam & Williams, 2022; Frandsen et al., 2020).

Studies in the field of evolutionary biology and ecological or conservation genetics of wild populations often focus on understudied organisms, for which there is rarely a chromosome-level reference genome and much less a known recombination map available. Even if a good reference genome is available, resequencing projects often need to use cost-effective sequencing approaches, like restriction site-associated DNA sequencing (RADseq) (Andrews et al., 2016), or low coverage whole genome sequencing (Lou et al., 2021), to maximize sample sizes at the expense of genome-wide coverage. While allowing larger sample sizes at an affordable cost, these sequencing techniques reduce the amount of information that can be extracted from the sequencing data. This often prevents the application of state-of-the-art methods developed for human or other model organisms. For example, it is usually not possible to model the positional auto-correlation of ancestry along the genome when only a subset of the genome is covered by sequencing reads, as in reduced representation sequencing, or when genotypes cannot be reliably called, as with low depth sequencing data. The same limitations apply in the absence of a highly contiguous reference genome where the sequencing data can be mapped to. In these situations, only the patterns of interancestry heterozygosity can be used to model the recent admixture history (Gompert et al., 2017).

Although there are several methods available to detect recent admixture or hybridization based on inferring genome-wide proportions of interancestry heterozygosities (Anderson & Thompson, 2002; Fitzpatrick, 2012; Crouch & Weale, 2012; Pfaffelhuber et al., 2022; Shastry et al., 2021; Wringe et al., 2017a, 2017b), all of them have at least one of two limitations. Most of these methods, with the exception of ENTROPY (Shastry et al., 2021), do not have an efficient implementation that can be applied to genome-scale data sets with large sample sizes. Moreover, only ENTROPY can be run from genotype likelihoods, allowing for analyses of low depth sequencing data. Additionally, most methods do not explicitly infer the underlying recent admixture pedigree and provide it as an output to the user. Only the Bayesian method NEWHYBRIDS (Anderson & Thompson, 2002) and associated packages such as *hybriddetective* (Wringe et al., 2017b) classify the samples in different hybrid classes (F1, F2, backcrosses…) and assign posterior probabilities for each class, hence quantifying the confidence in the classification. However, these methods require as input a pre-specified set of hybrid classes that can be detected, defined each by their expected values of interancestry heterozygosities. This will necessarily result in misclassification if the recent admixture history of a target sample is not included in the set of potential hybrid classes. Furthermore, it is limited to admixture between two parental populations (K=2). Admixture in nature is often complex and can potentially involve more than two populations. Alternative approaches are based on directly interpreting the estimates of interancestry heterozygosity patterns relative to their expectation for different hybrid classes (Pfaffelhuber et al., 2022; Shastry et al., 2021; Gompert et al., 2014; Waples et al., 2021). These can potentially be applied to situations with multi-population admixture. However, none of these methods output straightforward evaluations of recent admixture histories compatible with the estimates, or compare between the relative fit of different admixture histories. Therefore, there is a considerable unexploited potential in methods that make more explicit use of the information contained in interancestry heterozygosity for inferring concrete recent admixture histories in a quantitative and easily interpreted framework.

To address this situation we implement two similar, but complementary models that infer patterns of interancestry heterozygosity, and use the estimates to explore and evaluate concrete recent admixture histories. The ‘paired ancestries model’ (Gompert et al., 2014; Nøhr et al., 2021) models the global proportions of homozygous and heterozygous ancestries, without distinguishing between the two possible permutations of heterozygous ancestry (unordered paired ancestries). In contrast, the ‘parental admixture model’ (Crouch & Weale, 2012; Pfaffelhuber et al., 2022) allows to estimate ordered paired ancestries that can distinguish between the two permutations, which increases the resolution to detect cases of recent admixture based on the paired ancestries (Figure S1). Both models have been previously described in studies addressing recent admixture and hybridization (Crouch & Weale, 2012; Pfaffelhuber et al., 2022; Shastry et al., 2021; Gompert et al., 2014), but its potential to directly infer recent admixture pedigrees has not yet been explored. Moreover, our implementation can be run from called genotypes or genotype likelihood data in reasonable times for large data sets, and is furthermore the first implementation of the parental admixture model for genotype likelihood data. We have implemented both methods as extensions to the software NGSremix (Nøhr et al., 2021). Our method can be applied on genomic data with reduced genome contiguity or low depth sequencing (Figure 1A), and is therefore suitable for non-model organism analyses. Furthermore, we develop the independent software `apoh` (Admixture Pedigrees Of Hybrids) to automate downstream interpretation of the estimated proportions in the context of recent admixture. This is done by exploring different recent admixture pedigrees and their relative fit to the estimated proportions. In `apoh` we also provide two complementary indices that are descriptive of the model fit and the recent admixture process, respectively. We implement apoh both as a command-line tool and as a Graphical User Interface (GUI) that facilitates their use, and the interpretation of results, in applied cases.

**Figure 1:**
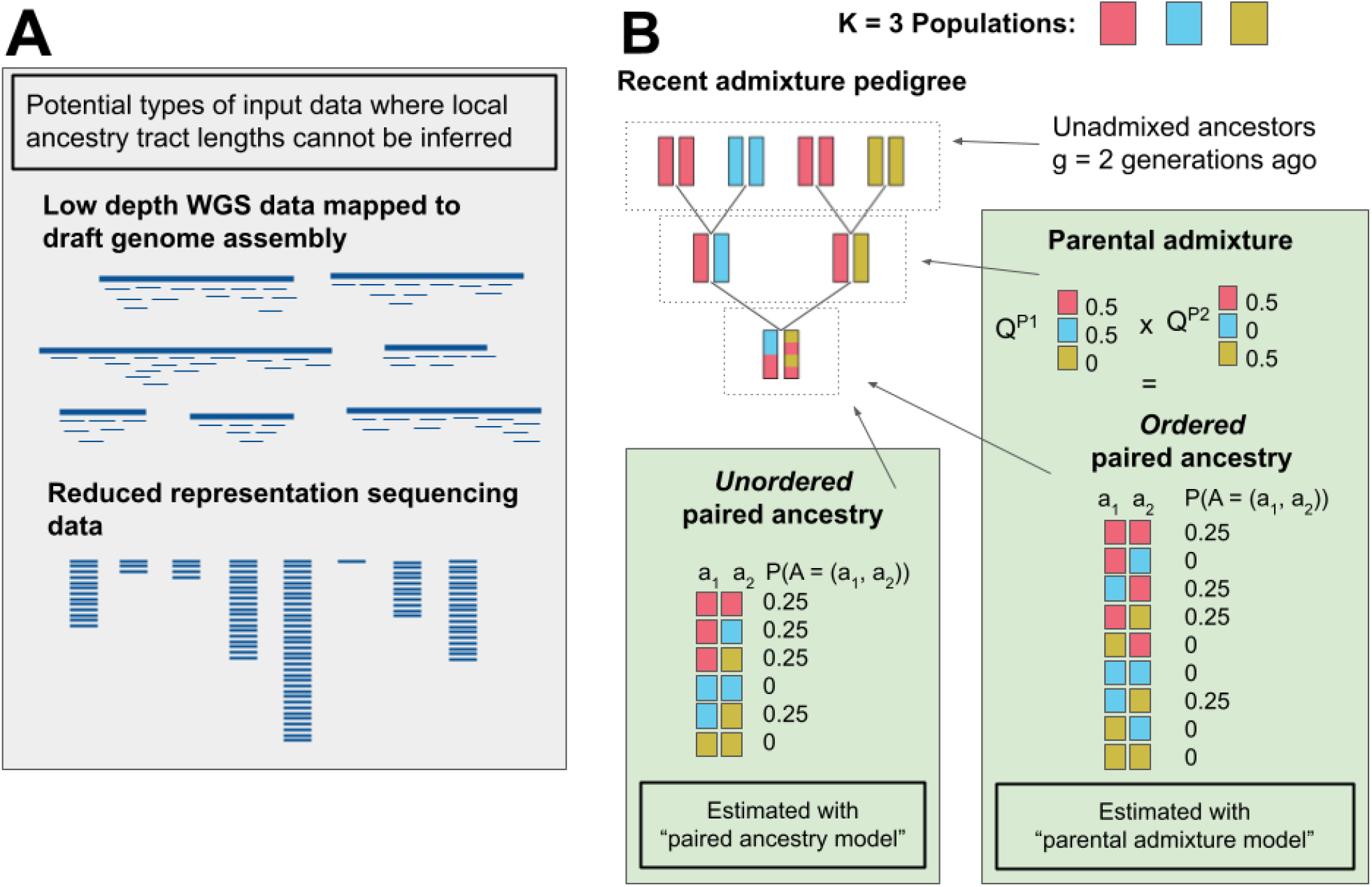
Schematic of the methods. A. Types of input data where the method can work, where other methods requiring genomics contiguity cannot be applied. B. Schematic of the two models of paired ancestry in the context of a recent admixture pedigree of an individual recently admixed between three different ancestries. Paired ancestries (left), ordered or unordered, are specific to the offspring, while parental ancestry (right) can be estimated from the offspring and in turn can be used to calculate ordered paired ancestry proportions.

## 2 Results

We present a method to evaluate a series of probable admixture pedigrees consistent with the estimated patterns of interancestry heterozygosities. The method works on a range of different data types, and notably both for low-depth and reduced representation sequencing data, including situations where such data is mapped to a draft genome assembly rather than a high quality chromosome level reference genome.

### 2.1 Models of paired ancestries

The inference of recent admixture in the methods presented here is based on modelling the patterns of interancestry heterozygosity, using the genome-wide proportions of all possible ancestry configurations across a large number of loci. Assuming a diploid genome, an admixed individual carries at each loci two alleles that can have either the same or different population ancestries. The possible ancestry pair configurations are then equal to all possible pairwise combinations of *K* ancestries. We refer to the vector *A* = (*a*_1_, *a*_2_), which specifies the population of origin of a pair of alleles at a locus, as the ‘*paired ancestry*’. The paired ancestries can be either ordered or unordered. The ordered version distinguishes the ancestry of the maternal and paternal allele, while in the unordered version the parental origin is ignored (Figure 1). We refer to the global genome-wide proportions of each paired ancestry as the ‘paired ancestry proportions’, which give the probability of observing a certain paired ancestry at a certain locus, *P*(*A* = (*a*_1_, *a*_2_)). The same or very similar quantities to the unordered paired ancestries have been used before in the study of recent admixture, with different names such as ‘genomic proportions’ (Fitzpatrick, 2012), ‘admixture class matrix’ (Gompert et al., 2014), ‘admixture complement matrix’ (Shastry et al., 2021) and ‘ternary ancestry fractions’ (Waples et al., 2021).

We use three different ancestry models to infer the global paired ancestry proportions. We give here a brief description of each model, the information they contain and their assumptions, while a more formal description can be found in the Materials and Methods section.

#### Standard admixture model

is the model of ancestry from the widely used software *structure* (Pritchard et al., 2000) and ADMIXTURE (Alexander et al., 2009). The ancestry of the pair of alleles at a locus is assumed to be independent given the individual’s admixture proportions *Q*, which is not the case in cases of recent hybridization. The estimated paired ancestries under this model, therefore, are consistent with the case where admixture and hybridization happened several generations ago such that *P*(*A* = (*a*_1_, *a*_2_)) ≈ *P*(*a*_1_)*P*(*a*_2_).

#### Paired ancestries model

explicitly infers the probability of each paired ancestry combination. It does not distinguish between different permutations of ancestry combinations, so *P*(*A* = (*a*_1_, *a*_2_)) = *P*(*A* = (*a*_2_, *a*_1_)). It therefore infers what we call ‘unordered paired ancestries’ (Figure 1B). It can model both an excess of heterozygous ancestries and an excess of homozygous ancestries, relative to the expected under the standard admixture model. This means it can account for cases of recent admixture, that will lead to an excess of heterozygous ancestries, and also for inbreeding, that leads to an excess of homozygous ancestry.

#### Parental admixture model

infers the admixture proportions of the two parents of the individual. The paired ancestry proportions are then given by the product of the parental admixture proportions. If the parents have different admixture proportions, the model will give different probability to each permutation of paired ancestries, *P*(*A* = (*a*_1_, *a*_2_)) ≠ *P*(*A* = (*a*_2_, *a*_1_)), so it infers ‘ordered paired ancestries’ (Figure 1B). It can account for cases of recent admixture, where the parents will have different admixture proportions and therefore there will be an excess of heterozygous ancestries. However, it cannot account for inbreeding, since it does not allow for excess of homozygous ancestry.

We implemented the ‘paired ancestry model’ and the ‘parental admxiture model’ as part of the NGSremix software (Nøhr et al., 2021). They can be used to estimate paired ancestry proportions and provide information on the recent admixture process (Figure 1). The paired ancestries model (Gompert et al., 2014; Nøhr et al., 2021) models the global proportions of homozygous and heterozygous ancestries, without distinguishing between the two possible permutations of heterozygous ancestry (unordered paired ancestries). In contrast, the parental admixture model (Crouch & Weale, 2012; Pfaffelhuber et al., 2022) allows to estimate ordered paired ancestries that can distinguish between the two permutations, which increases the resolution to detect cases of recent admixture based on the paired ancestries (Figure S1). The increase in resolution allows for improved identification of admixture events more than four generations ago, and distinguishes between different pedigrees. For this reason, we base most of the downstream inference on the ordered paired ancestries estimated with the parental admixture model. We did not implement the standard admixture model, since its parameters can be calculated as a function of the estimates from either of these two models, or inferred with established software such as ADMIXTURE for genotype data (Alexander et al., 2009) or NGSadmix for genotype likelihoods (Skotte et al., 2013), among others.

### 2.2 Inference of recent admixture pedigrees

We developed the software apoh to automatically explore the most compatible admixture pedigrees and assess if there is evidence for the sample being very recently admixed. From the ordered paired ancestry proportions estimated with the parental admixture model of an admixed individual, we explore the pedigrees that are most compatible with it assuming that either all ancestors where unadmixed four generations ago, or assuming an independent model where the ancestries of the two alleles are independent. Due to the inherent stochasticity in the parent-to-offspring transmission of haplotypes, we cannot always unambiguously identify a single admixture pedigree from the paired ancestry proportions, even if we had perfect estimates. Moreover, we want the method to work in situations where we have no information on genome location. In this situation we cannot model the local correlation of ancestry such as local ancestry tracts, which makes us unable to account for the statistical uncertainty associated with the inference. To reflect this and other potential sources of uncertainty, we choose instead to exhaustively explore and compare multiple similar pedigrees that are most compatible with the estimated paired ancestry proportions.

Finally, as a baseline to assess the support for recent admixture, we use a pedigree where all ancestors 4 generations ago have the same admixture proportions. In this case, the paired ancestry proportions expected under the standard admixture model should constitute a good model fit. Because of the assumption of independence in this model, we refer to this pedigree as the ‘independent pedigree’. We note, however, that this type of pedigree includes situations where both parents are in fact recently admixed, but have equal admixture proportions (e.g. Figure S1, F2 admixture). It is therefore important to keep in mind that in our framework, an inferred lack of support for recent admixture does not necessarily rule out that recent admixture did in fact take place.

### 2.3 Summary indices

As a guide to help in interpreting the model estimates and their fit to inferred recent admixture pedigrees, we developed several summary indices. The main index is designed to assess the compatibility of a potential admixture pedigree to the estimated paired ancestry proportions, by using the distance between the estimate and the paired ancestry proportions expected under that pedigree. We measure the distance as the Jensen - Shannon distance (JSD), which is a symmetrical measure of divergence between two probability distributions, to measure the distance between paired ancestry proportions. We use the ordered paired ancestries, since these have more information to distinguish between different admixture pedigrees (Figure S1). We then use the JSD to rank potential admixture pedigrees; this includes recent admixture pedigree but also the independent pedigree, which accounts for the case where there is not evidence of recent admixture.

We furthermore developed two additional summary indices that convey information contained in the estimates of paired ancestry proportions:

- The first one is the ‘inconsistency index’, which measures the distance between the unordered paired ancestries estimated with the paired ancestries model and the ones estimated with the parental admixture model. Since the two estimates are expected to be the same under a good model fit, a non-zero inconsistency index value suggests the estimates should be treated with caution. Non-zero distance can be due to inbreeding or to a bad model fit of the population allele frequencies used to model the sample ancestry.
- The second index is the ‘admixture index’, which, if the individual is in fact recently admixed, reflects the time since admixture. In the simplest case where the individual is the result of a single admixture event between two populations, the value of the index corresponds to the number of generations since admixture. In more complex cases, involving multiple admixture events and/or more than two ancestries, the index has a less straightforward interpretation, and is a lower bound on the sum of admixture times in the pedigree.

### 2.4 Implementation

We have implemented the parental admixture proportions model as an extension to the C++ software NGSremix (Nøhr et al., 2021), where we previously implemented the paired ancestry model for estimation of relatedness. It can be run either from genotype data or from genotype likelihood data. We implement the downstream inference of recent admixture pedigrees, visualization and derived summary indexes as an the software apoh, and developed a Graphical User Interface (GUI) using the ‘shiny’ R package (Chang et al., 2021). The GUI allows users to upload estimates of parental admixture proportions and paired ancestry proportions from NGSremix, and perform the downstream inference. All methods are publically available in github, NGSremix in https://github.com/KHanghoj/NGSremix and apoh in https://github.com/popgenDK/apoh.

### 2.5 Simulations

We first evaluated the performance of the parental admixture model under simple simulated scenarios of 100,000 unlinked markers. We simulated recently admixed individuals from two populations with varying number of generations since admixture and varying the amount of genetic differentiation between the two admixing populations. Ancestral allele frequencies were estimated from 20 unadmixed individuals from each ancestry together with 10 admixed individuals. A comparison of the true known minor parental ancestry with those estimated for the different admixture classes revealed that the inference is less accurate when the ancestral source differentiation is very low, and hence cannot distinguish between admixture classes when *F_ST_* is 0.01. However, at a still relatively minor *F_ST_* of 0.05 we are able to distinguish between different classes (Figure S2).

### 2.6 Evaluation using admixed family trios

To assess our implementation of the parental admixture proportions inference, we used genotype data of family trios from the 1000 Genomes project (Byrska-Bishop et al., 2022). Because the parents of the offspring were included in the unrelated samples for which we ran ADMIXTURE, we could compare direct estimates of the parent’s admixture proportions estimated with ADMIXTURE, and the estimates of these same proportions using only their offspring’s genotypes inferred by applying the parental admixture model. This revealed a high accuracy of the parental admixture model estimates (Figure 2). There are multiple cases where the two parents of an individual have different admixture proportions, which corresponds to a recent and ongoing admixture in these populations, as expected.

**Figure 2:**
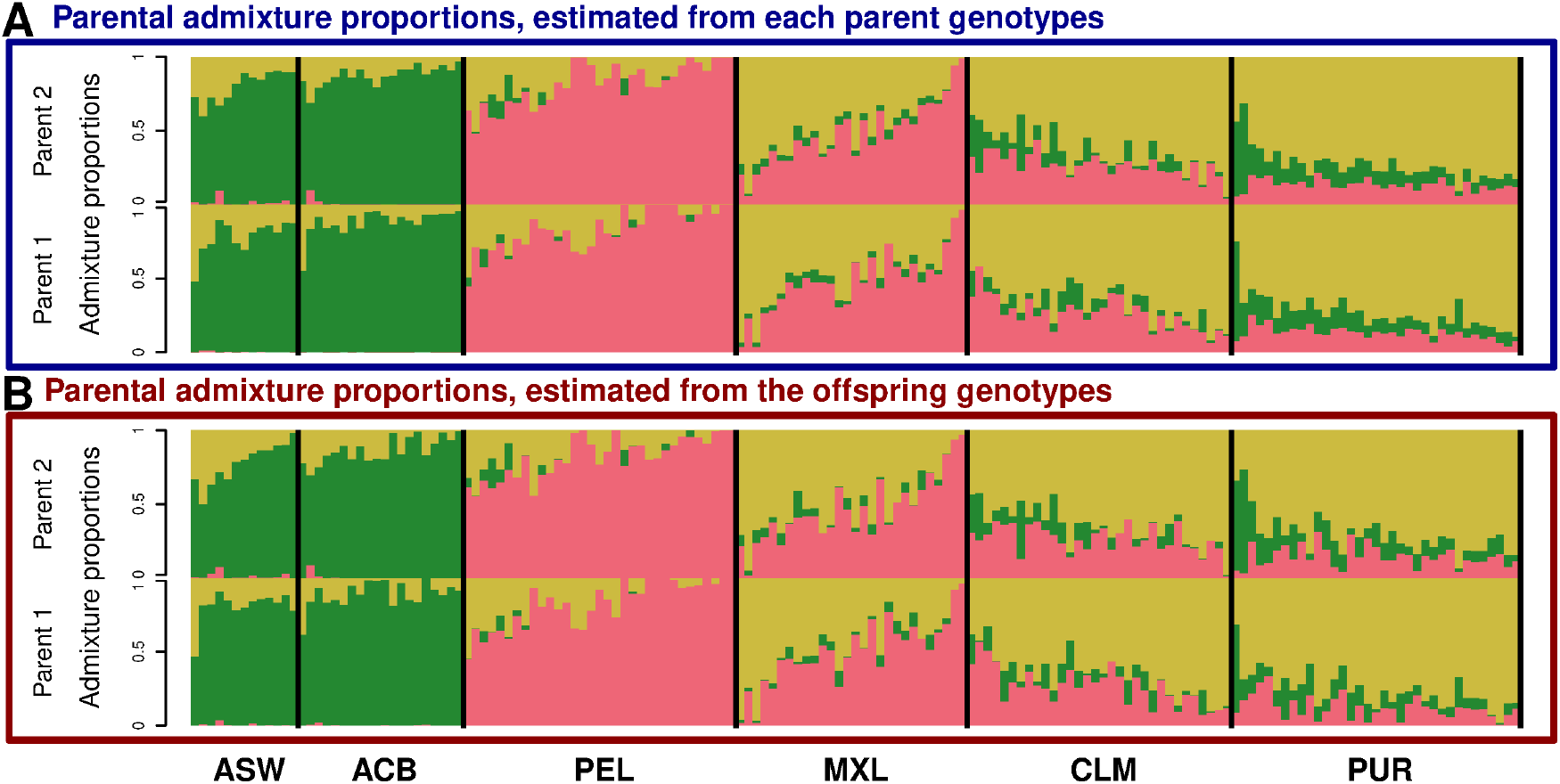
Estimates of parental admixture proportions from admixed 1000 genomes trios assuming *K* = 3 ancestral components. A. Admixture proportions estimated using ADMIXTURE from the parent’s genotypes. B. Admixture proportions estimated using the parental admixture model using only the child’s genotypes.

Furthermore, we also analyzed the offspring in trios from a South Asian population, PJL, in order to explore the performance of the method in cases of spurious admixture due to the source population not being well represented in the estimation of the population allele frequencies. We estimated paired ancestries with the parental and paired ancestry models for the PJL samples, using the population frequencies previously estimated with ADMIXTURE without including the PJL population. The parental admixture proportions modelled all PJL individuals as being admixed between all three ancestral populations that represent African, Native American and European ancestries (Figure S3). Therefore, these allele frequencies are not appropriate for modelling the South Asian ancestry of the PJL samples. Individuals from the PJL population show a relatively elevated inconsistency index compared to all other populations, demonstrating that the inconsistency index can detect violations of the model assumptions. An individual with sample ID HG01279 from the CLM population, for which the population allele frequencies are appropriate, show a relatively elevated inconsistency index (Figure S3). However, this individual has unusually high inbreeding relative to the other samples, demonstrating the other situation in which the inconsistency index value reflects departures from the model assumptions. Furthermore, five individuals from PJL with outlying inconsistency indices also showed high inbreeding coefficients, revealing that the inconsistency index reflects the cumulative effects of poorly estimated ancestral allele frequencies and inbreeding. We visualized the estimated unordered paired ancestry proportions for the HG01279 individual and for HG03707 from PJL, which has zero inbreeding and for which the high inconsistency index is therefore exclusively driven by poorly modelled allele frequencies. The plot illustrates that both of the model violations lead to an increase in homozygous paired ancestry proportions relative to the independent pedigree and the parental admixture model estimates (Figure S3).

### 2.7 Real data applications

To explore the applicability of the methods in studies with non-model organism presenting more challenging data, we used data sets from two African antelope species. Both of them included populations with recent gene flow between different species or sub-species. One data set is low-depth whole genome data (waterbuck), and the other is reduced representation sequencing (Grant’s gazelle). Therefore, these two data sets showcase the utility of our method for data typically used for non-model organisms where haplotype phasing or even genotype calling is not possible and where the position of SNPs on the chromosome is not known.

#### 2.7.1 Low depth WGS: waterbuck

We used low depth (3X) whole genome sequencing data for 38 waterbuck (*Kobus ellipsiprymnus*) samples (Wang et al., 2022) from four populations in East Africa. Two of the populations, Kidepo Valley National Park (KVNP, Uganda) and Maswa (Tanzania), belong to the defassa waterbuck subspecies (*K. e. defassa*). The other two populations, Samburu and Nairobi (Kenya), are in the range of the common waterbuck subspecies (*K. e. ellipsiprymnus*), but are known to have received recent gene flow from the defassa waterbuck (Figure 3A) (Wang et al., 2022; Lorenzen et al., 2006).

**Figure 3:**
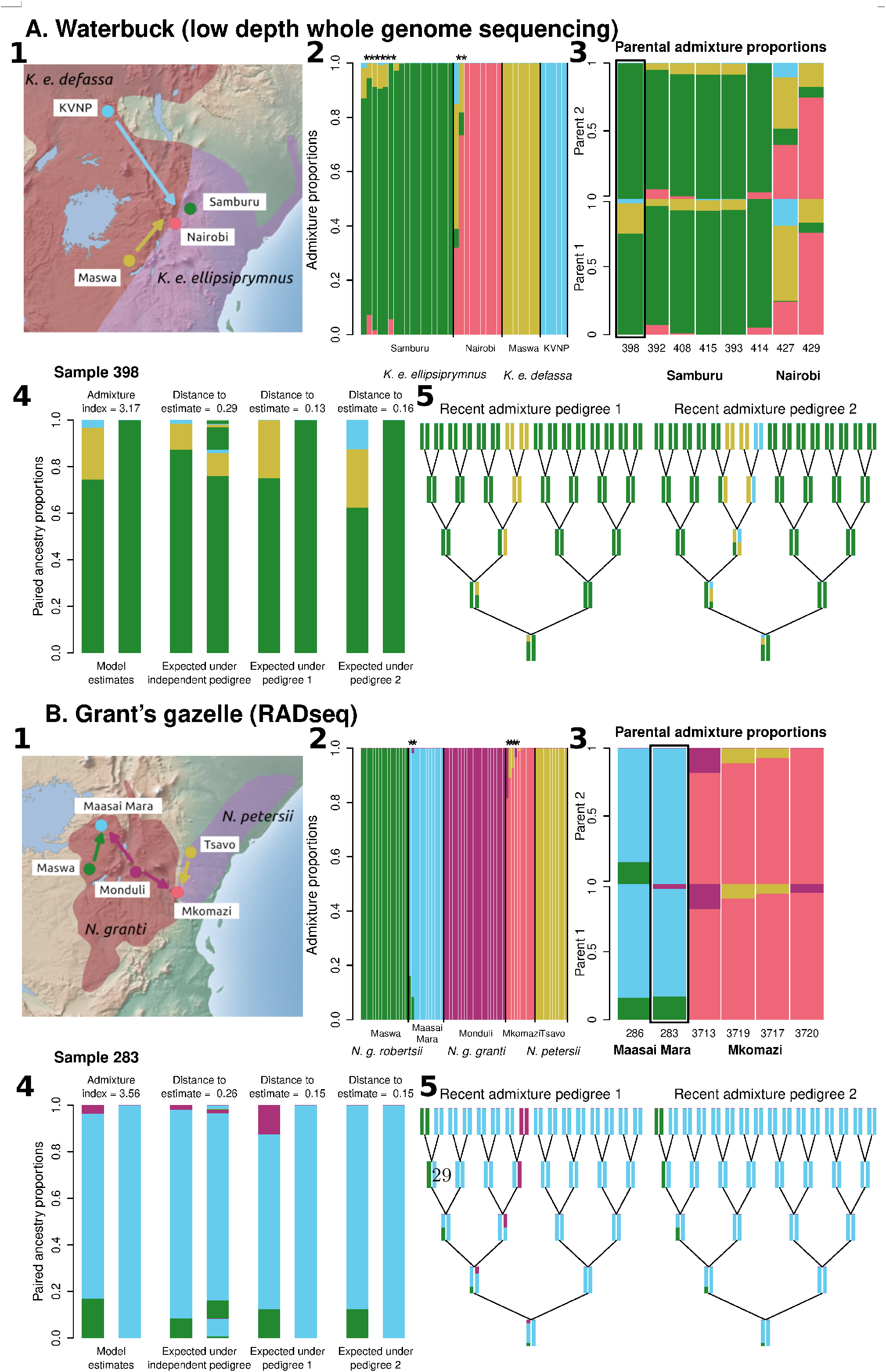
Application of the method to two datasets of non-model organisms with different sequencing strategies, A. low-depth whole genome sequencing data of waterbuck and B. RADseq data of Grant’s gazelle. For both A. and B., the upper panel shows: 1. A map of the sample localities with arrows representing gene flow movements (the origin of the arrows represents the most similar source populations, which are not necessarily the true source populations of the migrants). 2. A barplot of admixture proportions for all individuals, estimated with NGSremix, with admixed individuals for which we estimated paired ancestry proportions marked with an asterisk. 3. A barplot of parental admixture proportions for the admixed individuals, estimated using the parental admixture model. The recent admixture history as explored with apoh for the sample highlighted with a square is shown in the bottom panel. This panel shows: 4. The ordered paired ancestry proportions estimated with the parental admixture model, together with its admixture index, and the expected ordered paired ancestry proportions under independent pedigrees and under the two most compatible recent admixture pedigrees, with the distance to the model estimates for each of the three. 5. Visualization of the two most compatible recent admixture pedigrees.

Eight samples, six from Samburu and two from Nairobi, were modelled as having ancestry from more than two populations, all but one involving admixture between the two subspecies (Figure 3A). We ran our implementation of the paired ancestry and parental admixture models in these 8 samples, using the population allele frequencies estimated with NGSadmix at K=4, and estimated ordered and unordered paired ancestry proportions for all samples (Figure S4). Two of the samples, ID 398 from Samburu and 427 from Nairobi, had parents with different admixture proportions, which is the main signal of recent admixture (Figure 3A). We then applied apoh to infer the most compatible pedigrees and calculate the recent admixture indexes (Table 1). For sample 398 from Samburu, the distance to the most compatible recent admixture pedigree showed a clear improvement with respect to the independent pedigree. This, together with a low inconsistency index, indicates there is strong evidence that the individual is recently admixed and that the favored admixture pedigree is a good approximation. The most compatible recent admixture pedigree shows that the sample is a double backcross of a defassa/common waterbuck hybrid. Sample 427 from Nairobi shows a more complicated admixture history, with ancestry from four different populations (Figure S5). However this sample has a considerably higher inconsistency index, which means the paired ancestry estimates between the two models do not coincide (Table 1). This suggests the individual is probably recently admixed from a defassa waterbuck population that is not well represented in the data set, and therefore the allele frequencies are not accurate and prevent a reliable evaluation of its admixture pedigree.

**Table 1:** Summary indices for all admixed samples of waterbuck and Grant’s gazelle analysed. The inconsistency index indicates how well the ordered paired ancestries for which the pedigree inference is based fit the data. The distance to the different pedigrees is the Jensen-Shannon distance (JSD) between the expected paired ancestries under the pedigree and the paired ancestries estiamted by the model. The independent pedigree represents the case of not very recent admixture. The minimum distance is highlighted in bold for each sample. For those samples that a recent admixture pedigree fits the data better than the independent pedigree, the admixture index is shown, which indicates the time since admixture if there is a single admixture event (see Methods and Results for details).

| Species | SampleID | Inconsistency index | Distance to independent pedigree | Distance to pedigree 1 | Distance to pedigree 2 | Admixture index |
| --- | --- | --- | --- | --- | --- | --- |
| Waterbuck | 398 | 0.02 | 0.29 | **0.13** | 0.16 | 3.17 |
| Waterbuck | 392 | 0.1 | **0** | 0.25 | 0.28 |  |
| Waterbuck | 408 | 0.08 | **0.02** | 0.15 | 0.24 |  |
| Waterbuck | 415 | 0.07 | **0.04** | 0.11 | 0.22 |  |
| Waterbuck | 393 | 0.07 | **0.01** | 0.1 | 0.22 |  |
| Waterbuck | 414 | 0.05 | **0** | 0.16 | 0.19 |  |
| Waterbuck | 427 | 0.23 | 0.2 | **0.09** | 0.11 | 5.13 |
| Waterbuck | 429 | 0.15 | **0** | 0.12 | 0.14 |  |
| Grant’s gazelle | 286 | 0.41 | **0** | 0.06 | 0.1 |  |
| Grant’s gazelle | 283 | 0.01 | 0.26 | **0.15** | 0.15 | 3.56 |
| Grant’s gazelle | 3713 | 0.01 | **0** | 0.1 | 0.1 |  |
| Grant’s gazelle | 3719 | 0.19 | **0** | 0.03 | 0.24 |  |
| Grant’s gazelle | 3717 | 0.05 | **0** | 0.11 | 0.21 |  |
| Grant’s gazelle | 3720 | 0.01 | 0.13 | **0.09** | 0.19 | 3.93 |

The remaining samples showed no support for a recent admixture pedigree. This indicates that admixture happened more than five generations ago, and has potentially involved cross-mating between admixed individuals (Figure S4). Sample 429 from Nairobi has an elevated inconsistency index and a complex admixture proportions pattern involving three ancestries (Figure S4), as does the recently admixed 427 from the same locality. Together, these results suggest the inconsistency in this case is due to the recent gene flow into Nairobi being from a population not well represented in the data.

#### 2.7.2 RADseq: Grant’s gazelle

We also assessed the performance of the methods on reduced representation sequencing data by applying them to a RADseq dataset (Garcia-Erill et al., 2021). The Grant’s gazelle species complex (*Nanger sp*.) comprises three species with a non-overlapping distribution restricted to East Africa. Previous analyses of the data set revealed two contact zones between divergent lineages, one in the Maasai Mara (Kenya) between two *N. granti* subspecies, *N. g. granti* and *N. g. robertsii*, and another in Mkomazi (Tanzania), between the *N. granti* and *N. petersii* species. Both populations show signs of ancient and recent admixture, suggesting lineages in this species complex have diverged with possibly recurrent periods of isolation and migration, and are currently in a time of lineage connectivity (Garcia-Erill et al., 2021). In addition to the admixed populations, we included samples from Maswa (Tanzania), which belongs to the *N. g. robertsii* subspecies of *N. granti*, Monduli (Tanzania) that belongs to *N. g. granti* subspecies, and Tsavo (Kenya), which belongs to the *N. petersii* species (Figure 3B).

We identified six samples with ancestry from multiple populations; two from Maasai Mara and four from Mkomazi. We used the inferred population allele frequencies to estimate ordered and unordered paired ancestries with the two paired ancestry models for all six admixed samples (Figure S4). The results identified two samples, one from each population, with signs of being recently admixed. Sample 283 from Maasai Mara shows recent ancestry from both *N.granti* subspecies. Two potential recent admixture pedigrees have nearly identical distance to the estimated ordered paired ancestry proportions. The first pedigree includes an ancestor from each of the two minor ancestries four generations ago, while the second only includes an ancestor from *N.g.robertsii* and disregards the lowest *N.g.granti* ancestry component (Figure 3B). This suggests that, in this case, there is not enough information in the estimates to distinguish between two fundamentally different admixture histories. A potential explanation is that the recently admixed population source is not well represented in the data, and the minor ancestry from *N.g.granti* could therefore be an artifact. The inconsistency index does not detect it (Table 1, Figure S6), but it could be due to very low ancestry proportion from the potentially spurious source, *N.g.granti*. The other recently admixed sample is 3720 from Mkomazi, a historically admixed population in the contact zone between the *N.petersii* and *N.granti* species. This individual is modelled as a recent backcross admixed from an *N.g.granti* population. The JSD to the estimates show strong improvement of the most compatible pedigree with respect to both the independent pedigree and the second recent admixture pedigree, which shows the individual is likely a triple or quadruple backcross of a *N.granti* population (Figure S7), with its admixture index ~ 5 (Table 1) favoring a quadruple backcross.

The remaining samples did not show evidence of recent admixture. Sample 286 from Maasai Mara and sample 3719 from Mkomazi had an elevated inconsistency index (Table 1). This is likely due to inbreeding, since other individuals with similar ancestry profiles from the same populations show no increased inconsistency index (Figure S6), suggesting the ancestry sources are good approximations.

## 3 Discussion

We have developed methods and summary indices that use the information contained in the paired ancestry proportions to learn whether there is evidence for recent admixture, and to characterize the recent admixture pedigree in case there is. We furthermore implemented two previously described models (Crouch & Weale, 2012; Pfaffelhuber et al., 2022; Shastry et al., 2021; Nøhr et al., 2021) to estimate paired ancestry proportions. The use of two models with complementary information allows better utilization of the information contained in the data. Our methods and implementation have minimal requirements in terms of the genomic data used for the analyses, making them applicable to situations with reduced resource availability, such as low depth sequencing data, analyses based on draft reference genome assemblies or reduced representation sequencing of non-model organisms. At the same time, we present an efficient implementation of the models that allows analysis of genome scale data sets with many samples. We evaluate the parental admixture model using simulated data and data with genotype information from trios. In this way we are able to compare the inferred parental ancestry from the offspring to that estimated directly from the parents. We also show that our index for consistency was able to indicate violations of the model such as inbreeding and ghost admixture.

To illustrate potential usages of our methods we apply it to some very challenging data based on low depth WGS mapped to a draft assembly, and RADseq data where individuals showed signs of being admixed between two to four different populations. These case studies highlight how the wide applicability of the method can provide insights into biological questions. In this case, we combined analyses of two data sets of bovid species from the same geographical area in East Africa. In both cases our analyses provide the strongest evidence yet presented for recent gene flow between historically isolated lineages in these two species complexes, which has been suggested before (Lorenzen et al., 2006; Garcia-Erill et al., 2021; Arctander et al., 1996; Lorenzen et al., 2008), but never formally tested. Both cases include at least one case of confirmed recent gene flow through a geographical barrier, the Rift Valley, that has historically separated the *N. g. grant*i and *N. g. robertsi* subspecies in the Grant’s gazelle, and the *K. e. ellipsiprymnus* and *K. e. defassa* subspecies in the waterbuck. For these two species cases, we are able to identify plausible admixture pedigrees, which provides details regarding the first few generations of hybridization between the different lineages in each case. Such detailed insight into the recent admixture history involving two full or incipient species is extremely valuable both for answering fundamental evolutionary biology questions, e.g. related to the speciation process, but also for practical conservation of the involved taxa. With such information in hand we have tangible evidence concerning the recent migration of individuals in the particular geographical setting, as well as the patterns of backcrossing taking place the first few generations after the initial hybridization event. Even in the cases where our method did not allow for a detailed admixture pedigree inference, it still provides clues regarding which of the modeling assumptions are violated and thereby provides insights into the admixture history. For example, in both the Grant’s gazelles and waterbuck we identified several admixed individuals where the admixture probably took place > 4 generations ago, and individuals where the estimated admixture proportions are biased due to a lack of representation of their ancestry sources among the data.

Often there are several possible recent admixture pedigrees that can lead to similar paired ancestries as inferred from the data. To deal with this uncertainty, our method evaluates several plausible pedigrees and prioritizes them based on proposed ad-hoc indexes and JSD distance between the inferred paired ancestry and those of the candidate pedigrees. We chose not to implement formal statistical tests of the relative fit between pedigrees, as done in other implementations of the models (Pfaffelhuber et al., 2022), because we specifically aimed our method to be applicable for genome-scale data sets without a chromosome-level reference genome. In this situation, sites are not independent because of linkage disequilibrium (LD) and the absence of reliable genome locations makes it impossible to account for the correlation introduced by long admixture tracts in cases of recent admixture. It has been shown that the accuracy and resolution of recent admixture inference is increased when admixture tracts can be reliably identified, compared to methods based solely on patterns of interancestry heterozygosity, such as ours (Avadhanam & Williams, 2022). Therefore, our method is not intended to supplant or compete with methods based on inferred admixture tract lengths, but rather to make recent admixture history accessible for cases where the reference genome contiguity or the sequencing data precludes the application of such methods. Consequently, our method focuses on developing indices that can aid in identifying, ranking and prioritizing the most compatible recent admixture history, without explicit statistical testing of relative fit.

In conclusion, we present a simple and user-friendly set of methods to estimate patterns of paired ancestry proportions from genetic data. These methods facilitate the identification of recently admixed individuals and characterization of their admixture history, which is done through evaluation of recent admixture pedigrees. The methods carries/imposes minimal requirements for the amount of information present in the sequencing data, which makes them widely applicability in studies of non-model organisms with limited availability of genomic resources.

## 4 Methods

### 4.1 Models of paired ancestries

We use ‘paired ancestries’ to model the patterns of interancestry heterozygosity. A paired ancestry is a vector *A* = (*a*_1_, *a*_2_) that specifies the ancestry of each allele pair at a locus (for a diploid organism). The genome-wide paired ancestry proportions can be estimated with different models of ancestry. The models we use share the basic form of the standard admixture model from the widely used software ADMIXTURE and *structure* (Pritchard et al., 2000; Alexander et al., 2009), but they differ in how paired ancestries are modelled. For simplicity, we assume that the population allele frequencies *F* for each of the *K* populations are known. These can be obtained from ADMIXTURE if called genotypes are available, or from NGSadmix (Skotte et al., 2013), when working with low depth sequencing data. Neither method use information from the genomic position and can therefore work for RADseq data, or when working with draft genome assemblies as reference. We denote the genetic data for the focal individual as *X*, which can either be genotype data or genotype likelihoods for *M* sites. For both the independent ancestry model used in ADMIXTURE, the paired ancestry model and the parental ancestry model we can can write the likelihood as

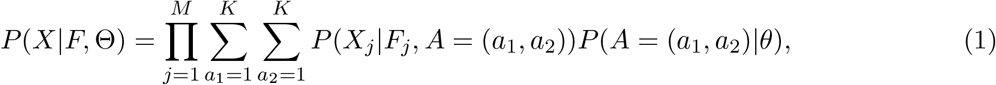

where *A* = (*a*_1_, *a*_2_) indicates the paired ancestries, and the parameters θ will vary depending on the parameterization of the paired ancestries in each model, which is described in the following sections. *F_j_* = (*f*_*j*1_, *f*_*j*2_,…, *f*_*jK*_) are the *K* ancestral population allele frequencies for site *j*. The calculation of *P*(*X_j_*|*A* = (*a*_1_, *a*_2_), *F_j_*) depends on whether we have called genotype data or genotype likelihood data. With genotype data *X* is an *M* length vector where *X_j_* ∈ {0, 1, 2}for site *j*, and we can calculate the probability as

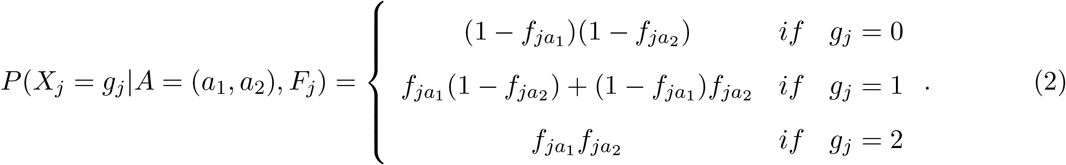

When working with genotype likelihoods, *X* is the sequencing data for *M* sites. In this case the probability of the data is given by

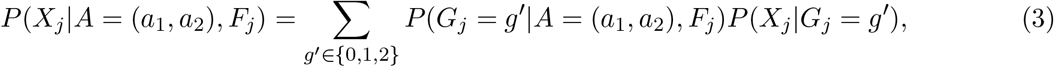

where *P*(*X_j_*|*G_j_* = *g*′) is the genotype likelihood for genotype *g*′ at site *j*. They can be estimated from aligned sequencing data with software such as GATK (McKenna et al., 2010), ANGSD (Korneliussen et al., 2014) or samtools (Li, 2011).

Different models of paired ancestries will differ in the parametetrization of *P*(*A* = (*a*_1_, *a*_2_)|*θ*). In the following, we briefly describe how paired ancestries are modelled in the standard admixture model and in the two explicit models of paired ancestries we implement.

#### 4.1.1 Standard Admixture model

In the standard admixture model (Pritchard et al., 2000; Alexander et al., 2009), *θ* corresponds to the admixture proportions *Q* = (*q*_1_, *q*_2_,…, *q_k_*). Under the admixture model, the population of origin of a pair of alleles is assumed to be independent given the admixture proportions, since

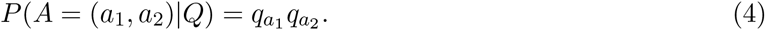

This assumption does not hold for inbreed individuals or individuals who are very recently admixed. Both of the other models that we use to infer recent admixture avoid this latter assumption, and use the patterns of paired ancestry to detect and characterize cases of recent admixture.

#### 4.1.2 Paired ancestry model

In the paired ancestry model (Gompert et al., 2014; Nøhr et al., 2021), we use a vector Φ containing 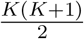 parameters, with Φ = (*ϕ*_11_, *ϕ*_12_,…, *ϕ*_*K*−1*K*_, *Φ_KK_*), where *ϕ*_*a*1*a*2_ denote the proportion of sites from the individual’s genome that carry one allele from population *a*_1_ and the other from population *a*_2_. The probability of a paired ancestry is therefore directly specified by Φ

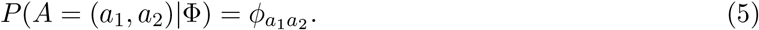

This model does not distinguish between the two possible ordering of paired ancestries, *P*(*A* = (*a*_1_, *a*_2_)) = *P*(*A* = (*a*_2_, *a*_1_)) = *ϕ*_*a*1*a*2_. It therefore only allows estimating ‘unordered paired ancestries’, as introduced above.

#### 4.1.3 Parental admixture model

In the parental admixture model (Crouch & Weale, 2012; Pfaffelhuber et al., 2022), we model the ancestry of the sample using its two parental admixture proportions. For *K* ancestral populations, we have 2*K* – 2 free parameters, with two *K* length vectors giving the admixture proportions of each parent, 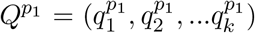 and 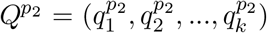 The probability of a paired ancestry proportion is then given by

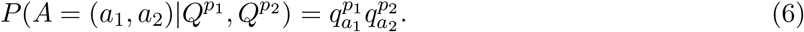

This model allows distinguishing the ordering of paired ancestries, since under this model *P*(*A* = (*a*_1_, *a*_2_)) ≠ *P*(*A* = (*a*_2_, *a*_1_)). Therefore, from this model we can calculate both unordered and ordered paired ancestries (Figure 1). Note that the method will have two equal solutions where the *P*_1_ and *P*_2_ labels are switched. For most downstream inference, we use the ordered paired ancestry estimates from this parental ancestry model, since its estimate relate directly to the recent admixture history of the individual.

### 4.2 Summarizing the information in the paired ancestry proportions

Given an estimate of the parental admixture proportions of an admixed individual, we can find the admixture pedigrees most compatible with the estimated proportions. We need to assume that the individual is the result of recent admixture, and that *g* generations back all its ancestors are homogeneous from one of the ancestries. We can define an admixture pedigree, going back *g* generations ago and involving *K* ancestries, as two *K* length vectors 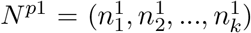 and 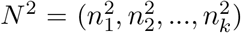. Each 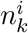 gives the number of ancestors of parent *i* = 1, 2 that are unadmixed from ancestry *k*, such that 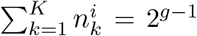. The parental admixture proportions have no information to distinguish the ordering of ancestors of each parent in the pedigree, so any permutation of the set of ancestors is equivalent. Because the space of potential recent admixture pedigrees under this conditions is relatively limited, we chose to explore all compatible recent admixture pedigrees. The only constrain is that we only consider recent admixture pedigrees with admixture proportions in the vicinity of the estimated parental admixture proportions. Furthermore, we also consider the ‘independent pedigree’, that corresponds to the pedigree where all ancestors have the same admixture proportions, such that it is compatible with a case of not so recent admixture.

Our approach of exploring and comparing multiple recent admixture pedigrees requires some way of ranking them by how well they explain the paired ancestry estimates. For this we use the Jensen-Shannon distance (JSD) between the ordered paired ancestry proportions estimated with the parental admixture model, and the ordered paired ancestry proportions expected under each of the considered admixture pedigrees. The JSD is a symmetrical measure of the distance between two probability distributions, and ranges from 0 to 1. In the context of comparing two paired ancestry proportions *P*_1_ and *P*_2_, the JSD is given by

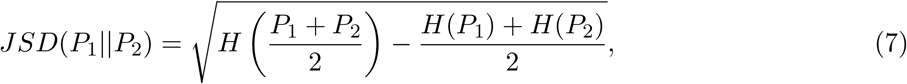

where *H*(*P_x_*) gives the entropy of the paired ancestry distribution, which in this context can be expressed as

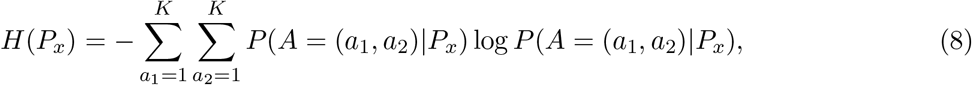

for *x* = 1, 2, for ordered paired ancestries, and

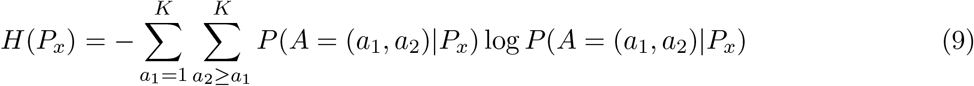

for *x* = 1, 2, for unordered paired ancestries.

In cases of complex admixture or when analysing many individuals, it can be difficult to interpret the results based on the paired ancestry proportions estimates and their fit to proposed admixture pedigrees. For this reason, we introduce two complementary indices that can be used as summary statistics to guide and prioritize the interpretation.

- **Inconsistency index**: this index is intended as an indicator of the reliability of the paired ancestry estimates. It is based on assessing the consistency between the paired ancestry estimates from the paired ancestry model, *P*_paired_, and the ones from the parental admixture model, *P*_parental_. We measure the consistency using the *JSD* between unordered paired ancestries probabilities defined by each model, such that a value of 0 indicates maximum consistency,

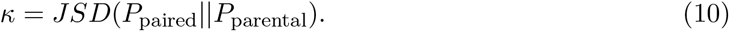 This index reflects violations of the two main model assumptions in the parental ancestry model. The first one is the the assumption of no inbreeding, which results in a higher proportion of homozygous paired ancestries than expected from a bifurcating pedigree. The paired ancestry model allows for excess of homozygous ancestries and thus does not make this assumption, which leads to the two models having different estimates in the case of inbreeding. The second assumption is that of having good estimates of allele frequencies for the populations from which the individual derives its ancestry. Many common situations, like admixture from ghost populations or continuous population structure, can lead to spurious admixture model results (Lawson et al., 2018; Garcia-Erill & Albrechtsen, 2020). While erroneous estimates of ancestral allele frequencies from such cases violate the assumptions of both models, it also results in different estimates between the two models. Therefore, the inconsistency index will also be elevated in the presence of this violation, making the index informative about this complication as well (see results).
- **Admixture index**: if an individual is recently admixed, the proportion of heterozygous ancestry contains information on the admixture time and the number of admixture events. This index aims to reflect this information and is defined as

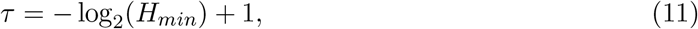

where *H_min_* is the lowest non-zero proportion of heterozygous ancestry. In the simplest case with only one recent admixture event between two ancestries, *τ* is equal to the admixture time in generations. In more complex cases involving multiple ancestries, if we assume each ancestry is different from the others and only admixes once, *τ* is the sum of all admixture times in generations, making it still informative on the admixture process. In any case, the index is a lower bound on the sum of admixture times. This index only makes sense if the individual is recently admixed and the estimated ordered paired ancestries are accurate. The value of *τ* therefore should not be considered unless the other metrics support this assumption, i.e. it is meaningless if the best fitting pedigree is the independent ancestries pedigree, or if an elevated inconsistency index suggests the ordered paired ancestry estimates are not accurate. For example, if an individual is admixed from an ancestry that is not well represented in our inferred ancestral population frequencies, the model would lead to estimates that would not be reflective of the true recent admixture history even if it is recently admixed.

### 4.3 Data analyses

#### 4.3.1 Simulations

We used the R package ‘bnpsd’ (Ochoa & Storey, 2021) to simulate allele frequencies for 100,000 independent sites in two populations with varying degrees of divergence. From these allele frequencies, we sampled genotypes for 20 unadmixed individuals for each of the two populations, and for 10 recently admixed individuals for each of 5 different recent admixture classes. Each class consisted of a single admixture event between the two populations, and varied by the number of generations that has passed since admixture, from one generation (resulting in a F1 admixed individual) up to five generations (quadruple backcross). For each simulation and admixture classes, we ran ADMIXTURE assuming K=2, and used the estimated population allele frequencies as input to estimate parental admixture proportions with our new implementation in NGSremix (Nøhr et al., 2021).

### 4.4 Admixed family trios from the 1000 Genomes Project

We tested our implementation of the parental admixture model using the family trios from the 1000G data set (Byrska-Bishop et al., 2022). We used the admixed populations from the Americas that have Native American ancestry: Peruvian in Lima Peru (PEL), Mexican Ancestry in Los Angeles, CA, USA (MXL), Colombian in Medellín, Colombia (CLM) and Puerto Rican in Puerto Rico (PUR), and the two groups with African ancestry, African Ancestry in SW USA (ASW), and African Caribbean in Barbados (ACB). We downloaded the publically available genotype calls for all autosomes, and used bcftools v. 1.10 (Danecek et al., 2021) to keep only diallelic SNPs. We then used plink v1.9 (Purcell et al., 2007) to remove related samples from these 6 admixed populations, and from the African population Yoruba in Ibadan, Nigeria (YRI) and the European population Utah residents with Northern and Western European ancestry (CEU). After filtering SNPs with minor allele frequency below 0.05, we kept 7,337,603 SNPs for 711 unrelated individuals. We used that data as input for ADMIXTURE assuming K=3 to estimate the individuals admixture proportions and population allele frequencies. We did 3 independent optimization runs and, after assessing convergence by observing similar log likelihoods across runs, selected the results with the maximum final log likelihood.

We then used our implementation of the parental admixture model to estimate parental admixture proportions for the offspring of the trios from admixed populations. Furthermore, we also estimated parental admixture proportions for the offspring of the South Asian population Punjabi in Lahore, Pakistan (PJL), which had not been included in the estimation of ancestral population allele frequencies with ADMIXTURE. The aim in this case is to explore the performance of the method in cases of spurious admixture due to individuals with incorrect population allele frequencies. To assess the relationship between inbreeding and the inconsistency index, we estimated inbreeding coefficients for the trio offspring for which we had estimated paired ancestry proportions. We used PCAngsd (Meisner & Albrechtsen, 2018) implementation of the ngsF maximum likelihood method to estimate inbreeding (Vieira et al., 2013; Meisner & Albrechtsen, 2019), using the individual allele frequencies estimated using 3 principal components to control for population structure.

### 4.5 Low depth whole genome sequencing dataset (waterbuck)

We got access to low coverage (3X) sequencing data of waterbucks (*Kobus ellipsiprymnus*) mapped to the defassa waterbuck (*K. e. defassa*) draft genome (DFW) and a list of quality controlled sites (Wang et al., 2022). From that we estimated genotype likelihoods with ANGSD (Korneliussen et al., 2014) using the GATK genotype likelihood model (McKenna et al., 2010). After keeping only sites with minor allele frequency 0.05, we had genotype likelihoods for 9,581,116 SNPs from 38 individuals, from which we estimated admixture proportions and ancestral population allele frequencies with NGSadmix (Skotte et al., 2013), assuming K = 4. We did 20 independent runs and, after assessing convergence, selected the run with the maximum final log likelihood. We then estimated paired ancestry proportions with the two models for admixed individuals, defined as those whose major ancestry proportion was below 97% in the NGSadmix results.

### 4.6 RADseq dataset (Grant’s gazelle)

We obtained the publicly available alignment files of Grant’s gazelle (*Nanger sps*.) RADseq data mapped to the domestic cow bosTau8 (*Bos taurus*) reference genome, and a list of quality controlled sites (Garcia-Erill et al., 2020). We estimated genotype likelihoods using ANGSD with the GATK model, for 70 individuals from five populations. We estimated genotype likelihoods for a total of 33,656 SNPs with minor allele frequency higher than 0.05 and used these as input in NGSadmix assuming K = 5, with 20 independent runs and selecting the run with the maximum final log likelihood. We then estimated paired ancestry proportions with the two models for admixed individuals, defined as those whose major ancestry proportion was below 0.97 in the NGSadmix results.

## 6 Data availability

No new data was generated as part of this study. The variant calls of the 1000G project data can be accessed in http://ftp.1000genomes.ebi.ac.uk/vol1/ftp/data_collections/1000G_2504_high_coverage/working/20201028_3202_phased/, and the raw sequencing data for the Grant’s gazelle RADseq dataset and the waterbuck low depth sequencing datasets can be accessed in SRA BioProject IDs PRJNA673069 and XXXXXXXX, respectively.

## 7 Acknowledgement

We thank Hans Redlef Siegismund, Laura Bertola, Malthe Sebro Rasmussen, Ida Moltke and all the members of the Population and Statistical Genetics group at the University of Copenhagen for their input in previous versions of the manuscript. We also thank Xi Wang and Mikkel Schubert for their contribution in processing the waterbuck data set. CW and AA are supported by the Independent Research Fund Denmark (grant number: 8021-00360B) and the University of Copenhagen through the Data+ initiative.

## Notes

### Competing Interest Statement

The authors have declared no competing interest.

## References

Feder, J. L., Egan, S. P., & Nosil, P. (2012). The genomics of speciation-with-gene-flow. Trends in genetics, 28(7), 342–350.

Edelman, N. B., & Mallet, J. (2021). Prevalence and adaptive impact of introgression. Annual Review of Genetics, 55, 265–283.

Pritchard, J. K., Stephens, M., & Donnelly, P. (2000). Inference of population structure using multi-locus genotype data. Genetics, 155(2), 945–959.

Vaha, J. P., & Primmer, C. R. (2006). Efficiency of model-based Bayesian methods for detecting hybrid individuals under different hybridization scenarios and with different numbers of loci. Mol. Ecol., 15(1), 63–72.

Buerkle, C. A. (2005). Maximum-likelihood estimation of a hybrid index based on molecular markers. Molecular Ecology Notes, 5(3), 684–687.

Beugin, M. P., Gayet, T., Pontier, D., Devillard, S., & Jombart, T. (2018). A fast likelihood solution to the genetic clustering problem. Methods Ecol Evol, 9(4), 1006–1016.

Lawson, D. J., van Dorp, L., & Falush, D. (2018). A tutorial on how not to over-interpret STRUCTURE and ADMIXTURE bar plots. Nat Commun, 9(1), 3258.

Boecklen, W. J., & Howard, D. J. (1997). Genetic analysis of hybrid zones: Numbers of markers and power of resolution. Ecology, 78(8), 2611–2616.

Anderson, E. C., & Thompson, E. A. (2002). A model-based method for identifying species hybrids using multilocus genetic data. Genetics, 160(3), 1217–1229.

Fitzpatrick, B. M. (2012). Estimating ancestry and heterozygosity of hybrids using molecular markers. BMC Evol. Biol., 12, 131.

Crouch, D. J., & Weale, M. E. (2012). Inferring separate parental admixture components in unknown DNA samples using autosomal SNPs. Eur J Hum Genet, 20(12), 1283–1289.

Pfaffelhuber, P., Sester-Huss, E., Baumdicker, F., Naue, J., Lutz-Bonengel, S., & Staubach, F. (2022). Inference of recent admixture using genotype data. Forensic Sci Int Genet, 56, 102593.

Shastry, V., Adams, P. E., Lindtke, D., Mandeville, E. G., Parchman, T. L., Gompert, Z., & Buerkle, C. A. (2021). Model-based genotype and ancestry estimation for potential hybrids with mixed-ploidy. Molecular ecology resources, 21(5), 1434–1451.

Zou, J. Y., Halperin, E., Burchard, E., & Sankararaman, S. (2015). Inferring parental genomic ancestries using pooled semi-Markov processes. Bioinformatics, 31(12), i190–196.

Pei, J., Zhang, Y., Nielsen, R., & Wu, Y. (2020). Inferring the ancestry of parents and grandparents from genetic data. PLoS computational biology, 16(8), e1008065.

Avadhanam, S., & Williams, A. L. (2022). Simultaneous inference of parental admixture proportions and admixture times from unphased local ancestry calls. Am J Hum Genet, 109(8), 1405–1420.

Frandsen, P., Fontsere, C., Nielsen, S. V., Hanghøj, K., Castejon-Fernandez, N., Lizano, E., Hughes, D., Hernandez-Rodriguez, J., Korneliussen, T. S., Carlsen, F., Siegismund, H. R., Mailund, T., Marques-Bonet, T., & Hvilsom, C. (2020). Targeted conservation genetics of the endangered chimpanzee. Heredity (Edinb), 125(1-2), 15–27.

Andrews, K. R., Good, J. M., Miller, M. R., Luikart, G., & Hohenlohe, P. A. (2016). Harnessing the power of radseq for ecological and evolutionary genomics. Nature Reviews Genetics, 17(2), 81–92.

Lou, R. N., Jacobs, A., Wilder, A. P., & Therkildsen, N. O. (2021). A beginner’s guide to low-coverage whole genome sequencing for population genomics. Molecular Ecology, 30(23), 5966–5993.

Gompert, Z., Mandeville, E. G., & Buerkle, C. A. (2017). Analysis of population genomic data from hybrid zones. Annual Review of Ecology, Evolution, and Systematics, 48, 207–229.

Wringe, B. F., Stanley, R. R. E., Jeffery, N. W., Anderson, E. C., & Bradbury, I. R. (2017a). Parallelnewhybrid: An r package for the parallelization of hybrid detection using newhybrids. Molecular ecology resources, 17(1), 91–95.

Wringe, B. F., Stanley, R. R. E., Jeffery, N. W., Anderson, E. C., & Bradbury, I. R. (2017b). Hybrid-detective: A workflow and package to facilitate the detection of hybridization using genomic data in r. Molecular Ecology Resources, 17(6), e275–e284.

Gompert, Z., Lucas, L. K., Buerkle, C. A., Forister, M. L., Fordyce, J. A., & Nice, C. C. (2014). Admixture and the organization of genetic diversity in a butterfly species complex revealed through common and rare genetic variants. Molecular ecology, 23(18), 4555–4573.

Waples, R. K., Hauptmann, A. L., Seiding, I., Jørsboe, E., Jørgensen, M. E., Grarup, N., Andersen, M. K., Larsen, C. V. L., Bjerregaard, P., Hellenthal, G., Hansen, T., Albrechtsen, A., & Moltke, I. (2021). The genetic history of Greenlandic-European contact. Curr Biol, 31(10), 2214–2219.

Nøhr, A. K., Hanghøj, K., Garcia-Erill, G., Li, Z., Moltke, I., & Albrechtsen, A. (2021). NGSremix: a software tool for estimating pairwise relatedness between admixed individuals from next-generation sequencing data. G3 (Bethesda), 11(8).

Alexander, D. H., Novembre, J., & Lange, K. (2009). Fast model-based estimation of ancestry in unrelated individuals. Genome Res., 19(9), 1655–1664.

Skotte, L., Korneliussen, T. S., & Albrechtsen, A. (2013). Estimating individual admixture proportions from next generation sequencing data. Genetics, 195(3), 693–702.

Chang, W., Cheng, J., Allaire, J., Sievert, C., Schloerke, B., Xie, Y., Allen, J., McPherson, J., Dipert, A., & Borges, B. (2021). Shiny: Web application framework for r [R package version 1.7.1].

Byrska-Bishop, M., Evani, U., Zhao, X., Basile, A., Abel, H., Regier, A., Corvelo, A., Clarke, W., Musunuri, R., Nagulapalli, K., Fairley, S., Runnels, A., Winterkorn, L., Lowy, E., Consortium, H. G. S. V., Flicek, P., Germer, S., Brand, H., Hall, I.., … Zody, M. (2022). High-coverage whole-genome sequencing of the expanded 1000 genomes project cohort including 602 trios. Cell, 185(18), 3426–3440.

Wang, X., Pedersen, C. E. T., Athanasiadis, G., Garcia-Erill, G., Hanghøj, K., Bertola, L. D., Rasmussen, M. S., Schubert, M., Liu, X., Li, Z., Lin, L., Jørsboe, E., Nursyifa, C., Liu, S., Muwanika, V., Masembe, C., Chen, L., Wang, W., Moltke, I., … Heller, R. (2022). Persistent gene flow suggests an absence of reproductive isolation in an african antelope speciation model. bioRxiv.

Lorenzen, E. D., Simonsen, B. T., Kat, P. W., Arctander, P., & Siegismund, H. R. (2006). Hybridization between subspecies of waterbuck (Kobus ellipsiprymnus) in zones of overlap with limited introgression. Mol Ecol, 15(12), 3787–3799.

Garcia-Erill, G., Kjaer, M. M., Albrechtsen, A., Siegismund, H. R., & Heller, R. (2021). Vicariance followed by secondary gene flow in a young gazelle species complex. Mol Ecol, 30(2), 528–544.

Arctander, P., Kat, P. W., Aman, R. A., & Siegismund, H. R. (1996). Extreme genetic differences among populations of Gazella granti, Grant’s gazelle in Kenya. Heredity (Edinb), 76 (Pt 5), 465–475.

Lorenzen, E. D., Arctander, P., & Siegismund, H. R. (2008). Three reciprocally monophyletic mtdna lineages elucidate the taxonomic status of grant’s gazelles. Conservation Genetics, 9(3), 593.

McKenna, A., Hanna, M., Banks, E., Sivachenko, A., Cibulskis, K., Kernytsky, A., Garimella, K., Altshuler, D., Gabriel, S., Daly, M., & DePristo, M. A. (2010). The Genome Analysis Toolkit: a MapReduce framework for analyzing next-generation DNA sequencing data. Genome Res., 20(9), 1297–1303.

Korneliussen, T. S., Albrechtsen, A., & Nielsen, R. (2014). ANGSD: Analysis of Next Generation Sequencing Data. BMC Bioinformatics, 15, 356.

Li, H. (2011). A statistical framework for SNP calling, mutation discovery, association mapping and population genetical parameter estimation from sequencing data. Bioinformatics, 27(21), 2987–2993.

Garcia-Erill, G., & Albrechtsen, A. (2020). Evaluation of model fit of inferred admixture proportions. Molecular ecology resources, 20(4), 936–949.

Ochoa, A., & Storey, J. D. (2021). Estimating FST and kinship for arbitrary population structures. PLoS Genet, 17(1), e1009241.

Danecek, P., Bonfield, J. K., Liddle, J., Marshall, J., Ohan, V., Pollard, M. O., Whitwham, A., Keane, T., McCarthy, S. A., Davies, R. M., & Li, H. (2021). Twelve years of SAMtools and BCFtools. Gigascience, 10(2).

Purcell, S., Neale, B., Todd-Brown, K., Thomas, L., Ferreira, M. A., Bender, D., Maller, J., Sklar, P., de Bakker, P. I., Daly, M. J., & Sham, P. C. (2007). PLINK: a tool set for whole-genome association and population-based linkage analyses. Am. J. Hum. Genet., 81(3), 559–575.

Meisner, J., & Albrechtsen, A. (2018). Inferring Population Structure and Admixture Proportions in Low-Depth NGS Data. Genetics, 210(2), 719–731.

Vieira, F. G., Fumagalli, M., Albrechtsen, A., & Nielsen, R. (2013). Estimating inbreeding coefficients from NGS data: Impact on genotype calling and allele frequency estimation. Genome Res, 23(11), 1852–1861.

Meisner, J., & Albrechtsen, A. (2019). Testing for Hardy-Weinberg equilibrium in structured populations using genotype or low-depth next generation sequencing data. Mol Ecol Resour, 19(5), 1144–1152.

Garcia-Erill, G., Kjær, M., Albrechtsen, A., Siegismund, H., & Heller, R. (2020). Mapped read data and files and scripts from: Vicariance followed by secondary gene flow in a young gazelle species complex (Dataset) [https://doi.org/10.5061/dryad.pzgmsbcjn]. Dryad.

