## Supplementary material for "Estimating admixture pedigrees of recent hybrids without a contiguous reference genome"

#### 1 Appendix 1: Inference for models of paired ancestry

##### S1 Paired ancestries model: EM for $\Phi$

In the following $\Phi$ and $X$ refer respectively to the paired ancestry proportions and the sequencing data for a single individual. We use an expectation maximization (EM) algorithm to obtain a maximum likelihood estimate $\Phi$. We denote the $i$th iteration of the estimated pairwise ancestry proportion $\Phi^{(i)}$. Based on this estimate, the expectation of the total number of sites with paired ancestries $A = (a_1, a_2)$ is given by

$$\mathbb{E}[A = (a_1, a_2)|X, F, \Phi^{(i)}] = \sum_{j=1}^{M} P(A_j = (a_1, a_2)|X_j, F_j, \Phi^{(i)}) = \sum_{j=1}^{M} \frac{P(X_j|A_j = (a_1, a_2), F_j)P(A_j = (a_1, a_2)|\Phi^{(i)})}{\sum_{a_1'=1}^{K}\sum_{a_2'=a_1'}^{K} P(X_j|A = (a_1', a_2'), F_j)P(A = (a_1', a_2')|\Phi^{(i)})}. \tag{S1}$$

Then we can update the estimate of $\Phi$ by calculating each of its entries as

$$\phi_{a_1 a_2}^{(i+1)} = \frac{\mathbb{E}[A = (a_1, a_2)|X, F, \Phi^{(i)}]}{\sum_{a_1'=1}^{K} \sum_{a_2'=a_1'}^{K} \mathbb{E}[A = (a_1', a_2')|X, F, \Phi^{(i)}]}. \tag{S2}$$

Finally, we calculate the new expectation given $\Phi^{(i+1)}$ with equation (S1) and use the expectation to calculate a new estimate with equation (S2) iterating until convergence. A proof that this is an EM for a similar case can by found in **Skotte2013**.

#### S2 Parental admixture model: EM for $Q^{p_1}$ and $Q^{p_2}$

We can estimate $Q^{p_1}$ and $Q^{p_2}$ from genotype or genotye likelihoods with an EM similar to the one used for $\Phi$. An important difference however is that now sites with heterozygous ancestry are not exchangeable, since we are assigning each allele to each parent. This means we estimate the expectation of the total number of sites where allele j will have ancestry $a_i$ for $j \in (1, 2)$ and $i \in (1, 2...K)$. For example the expected number of sites where allele $a_1$ will have ancestry from population 1 is given by

$$\begin{aligned}\mathbb{E}[A = (a_1, a_2')|X, F, Q^{p_1(i)}, Q^{p_2(i)}] = \sum_{j=1}^{M} \sum_{a_2'=1}^{K} P(A = (a_1, a_2')|X_j, F_j, Q^{p_1(i)}, Q^{p_2(i)}) = \\ \sum_{j=1}^{M} \frac{\sum_{a_2'=1}^{K} P(X_j|A_j = (a_1, a_2'), F_j) P(A_j = (a_1, a_2'|Q^{p_1(i)}, Q^{p_2(i)})}{\sum_{a_1'=1}^{K} \sum_{a_2'=a_1'}^{K} P(X_j|A = (a_1', a_2'), F_j) P(A = (a_1', a_2')|Q^{p_1(i)}, Q^{p_2(i)})}.\end{aligned} \tag{S3}$$

Then we can update the estimates of $Q^{p_2}$ and $Q^{p_2}$ by calculating each of its entries as

$$q_{a_1}^{p_1(i+1)} = \frac{\sum_{a_1'=1}^{K} \mathbb{E}[A = (a_1, a_2')|X, F, Q^{p_1(i)}, Q^{p_2(i)}]}{\sum_{a_1'=1}^{K} \sum_{a_2'=1}^{K} \mathbb{E}[A = (a_1', a_2')|X, F, Q^{p_1(i)}, Q^{p_2(i)}]} \tag{S4}$$

and

$$q_{a_2}^{p_2(i+1)} = \frac{\sum_{a_1'=1}^{K} \mathbb{E}[A = (a_1', a_2)|X, F, Q^{p_1(i)}, Q^{p_2(i)}]}{\sum_{a_1'=1}^{K} \sum_{a_2'=1}^{K} \mathbb{E}[A = (a_1', a_2')|X, F, Q^{p_1(i)}, Q^{p_2(i)}]} \tag{S5}$$

### 2 Supplementary figures

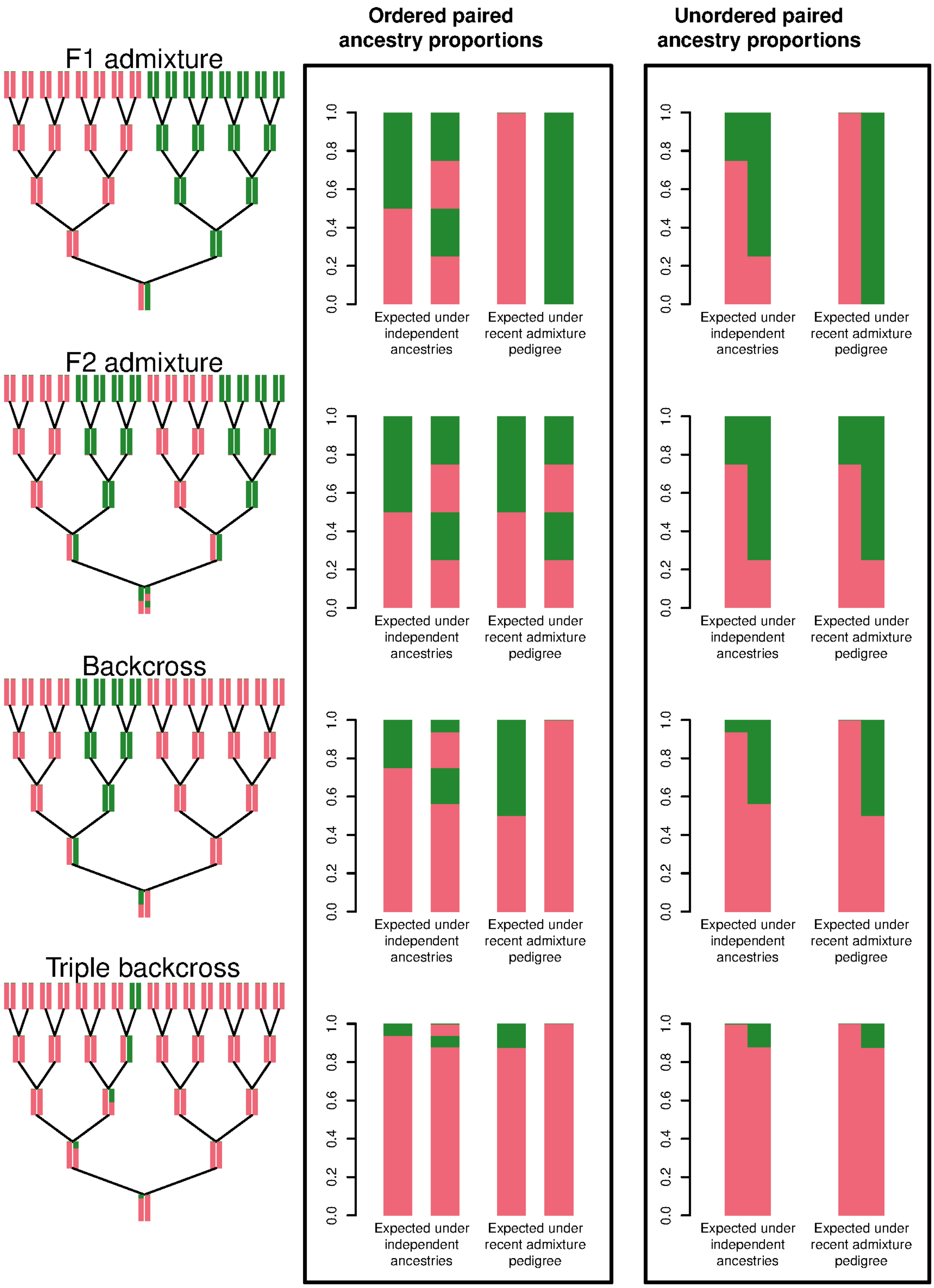


Figure S1: Pedigrees and expected paired ancestries for different recent admixture classes. For each, the left panel shows the recent admixture pedigree. The middle and right panel panel shows the ordered and unordered paired ancestry proportions, respectively, expected under an independent pedigree with same admixture proportions, and under the recent admixture pedigree.

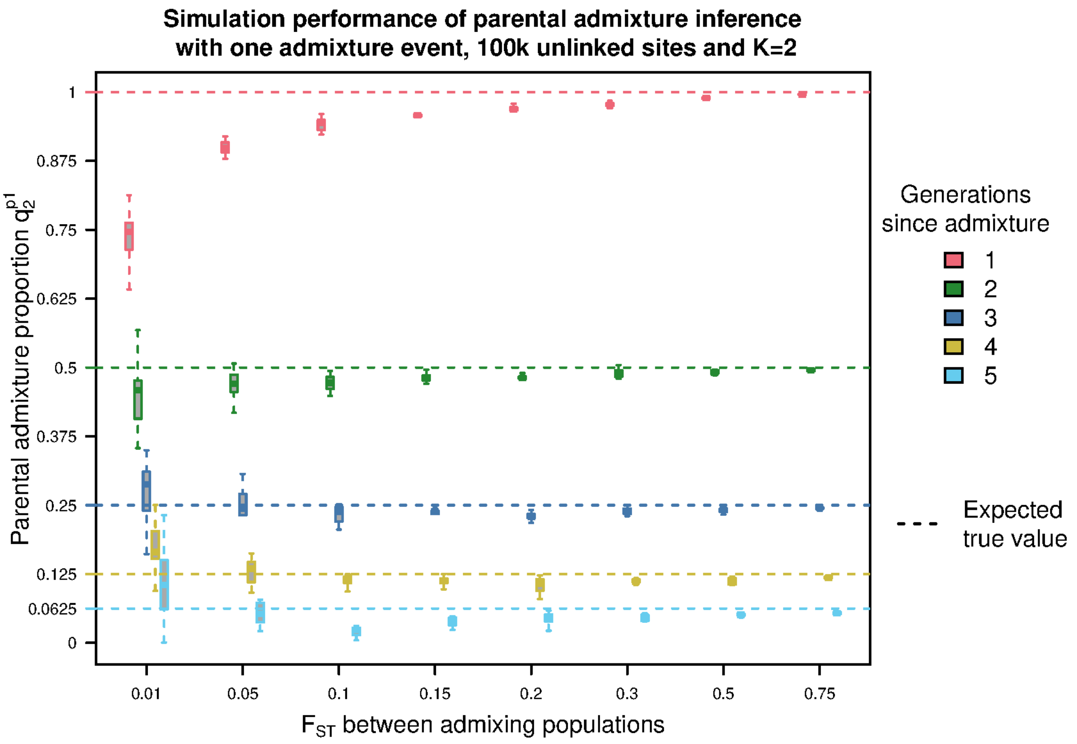


Figure S2: Evaluation of the parental admixture model inference accuracy using simulations. Admixture in all cases is a single admixture event between two different populations, and the recent admixture classes differ in the number of generations passed since admixture. The dashed lines indicate the parental admixture proportion value used in the simulation. For each admixture class and $F_{ST}$, the estimated parental admixture proportions for ten different replicates are plotted to measure the accuracy.

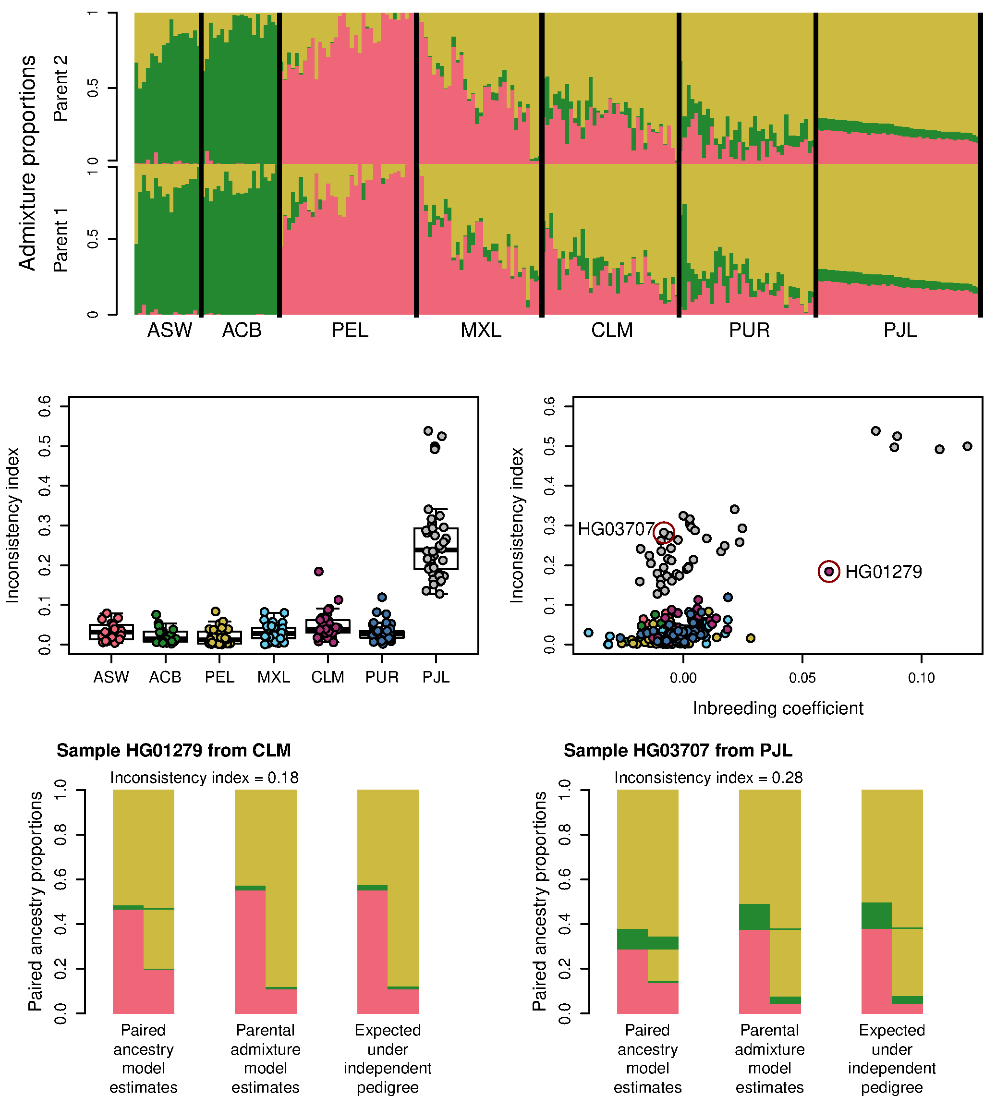


Figure S3: Evaluation of the inconsistency index with the 1000 genomes data. The top panel shows the parental admixture proportions estimated with the parental admixture model for all offsprings of family trios from the analyzed populations. The middle panels shows, on the left, the individual inconsistency index for all individual, separated by population, and the right these inconsistency indices are plotted agains estiamted inbreeding coefficient for each individual. The bottom panel shows, for two representative samples with a high inconsistency index due to inbreeding (left) and to a bad model fit (right), of the the estimated unordered paired ancestry proportions estimated under each model, whose distance give the inconsistency index, and expected under an independent pedigree.

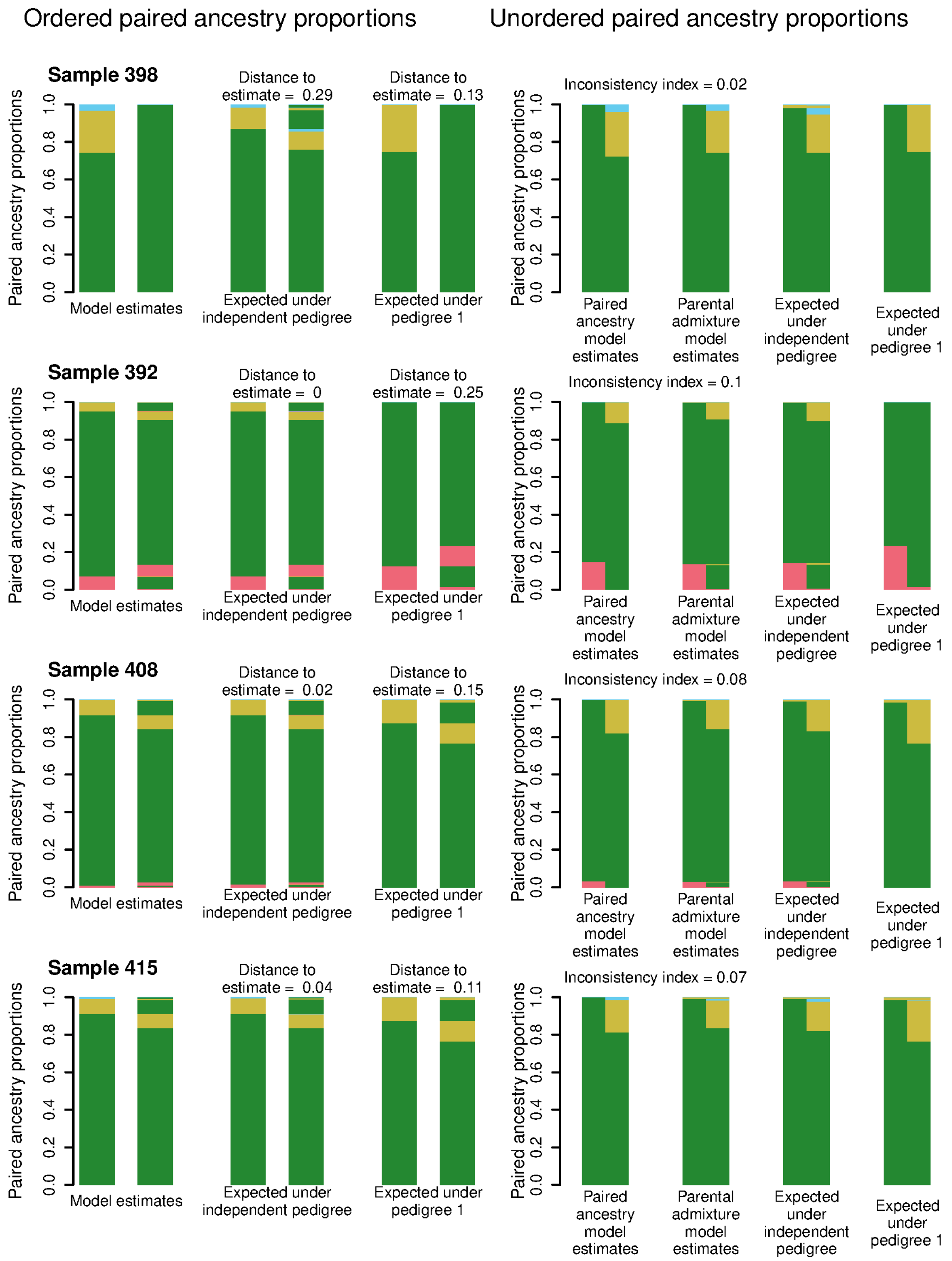


Figure S4: Paired ancestry proportions for all admixed waterbuck samples. Each row corresponds to a sample, where the left panel shows the ordered paired ancestries estimated with the parental admxiture model, the ones expected under the independent pedigree and the ones expected under the most compatible recent admixture pedigree. Distances for the pedigrees are the Jensen-Shannon distance (JSD) to the model estimates. The right panel shows the unordered paired ancestry proportions estimated with the paired ancestry model and the ones with the parental admixture model, the ones expected under an independent pedigree and the ones expected under the most compatible recent admixutre pedigree.

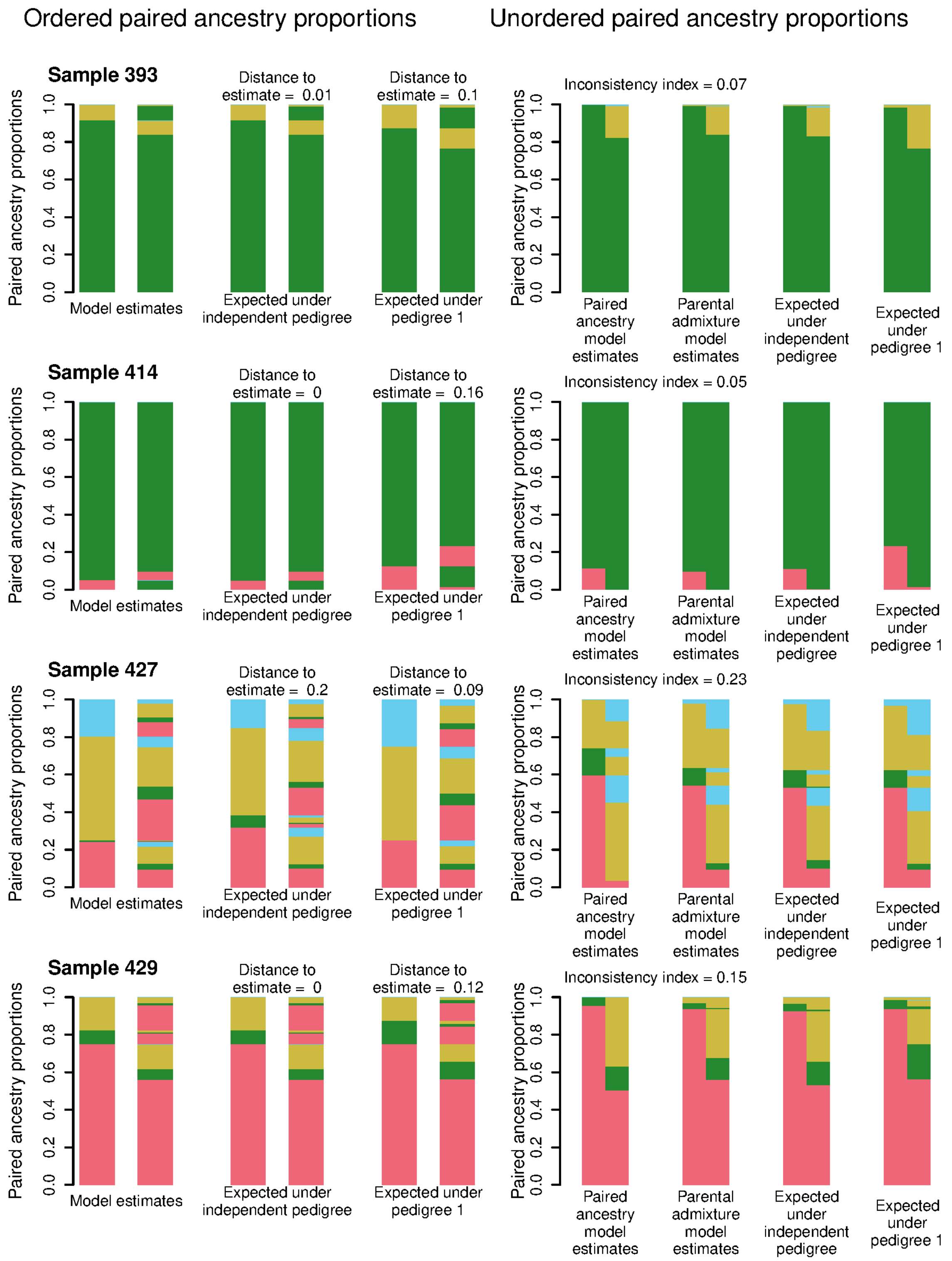


Figure S4: (continued) Paired ancestry proportions for all admixed waterbuck samples. Each row corresponds to a sample, where the left panel shows the ordered paired ancestries estimated with the parental admxiture model, the ones expected under the independent pedigree and the ones expected under the most compatible recent admixture pedigree. Distances for the pedigrees are the Jensen-Shannon distance (JSD) to the model estimates. The right panel shows the unordered paired ancestry proportions estimated with the paired ancestry model and the ones with the parental admixture model, the ones expected under an independent pedigree and the ones expected under the most compatible recent admixutre pedigree.

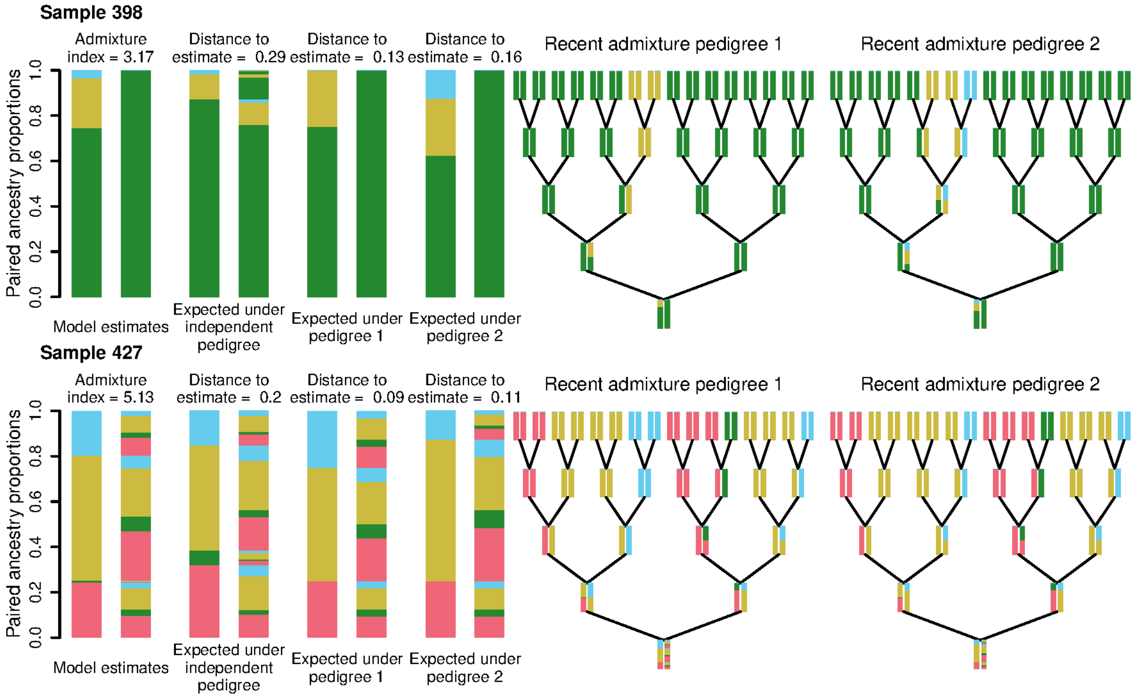


Figure S5: Ordered paired ancestry proportions and recent admixture pedigrees for the two waterbuck samples with evidence of being recently admixed. Each row corresponds to a sample, where the left panel shows the ordered paired ancestries estimated with the parental admixture model, the ones expected under the independent pedigree and the ones expected under the two most compatible recent admixture pedigrees. Distances for the pedigrees are the Jensen-Shannon distance (JSD) to the model estimates. The right panel shows the two most compatible recent admixture pedigrees.

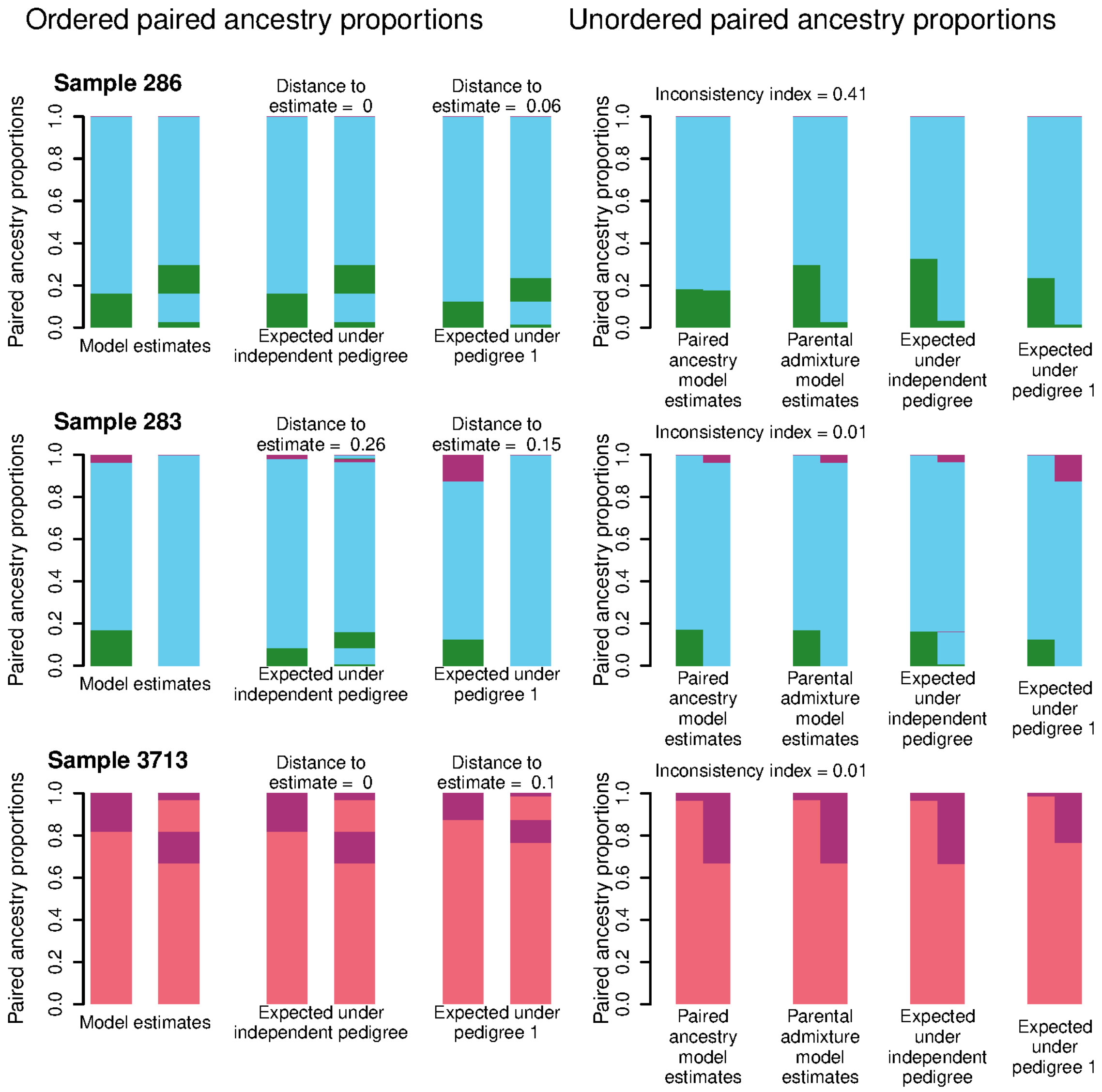


Figure S6: Paired ancestry proportions for all admixed Grant's gazelle samples. Each row corresponds to a sample, where the left panel shows the ordered paired ancestries estimated with the parental admxiture model, the ones expected under the independent pedigree and the ones expected under the most compatible recent admixture pedigree. Distances for the pedigrees are the Jensen-Shannon distance (JSD) to the model estimates. The right panel shows the unordered paired ancestry proportions estimated with the paired ancestry model and the ones with the parental admixture model, the ones expected under an independent pedigree and the ones expected under the most compatible recent admixutre pedigree.

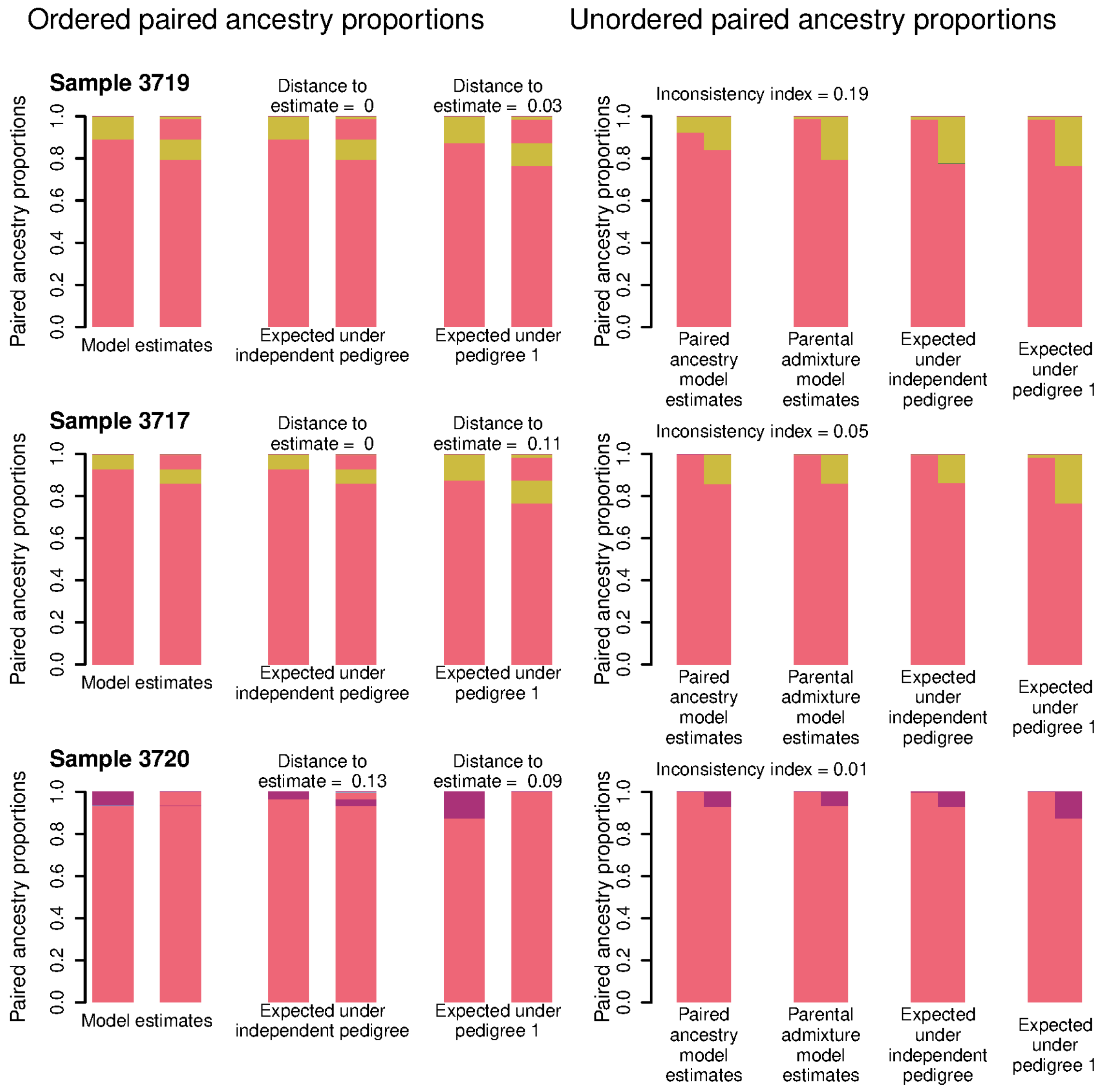


Figure S6: (continued) Paired ancestry proportions for all admixed Grant's gazelle samples. Each row corresponds to a sample, where the left panel shows the ordered paired ancestries estimated with the parental admxiture model, the ones expected under the independent pedigree and the ones expected under the most compatible recent admixture pedigree. Distances for the pedigrees are the Jensen-Shannon distance (JSD) to the model estimates. The right panel shows the unordered paired ancestry proportions estimated with the paired ancestry model and the ones with the parental admixture model, the ones expected under an independent pedigree and the ones expected under the most compatible recent admixutre pedigree.

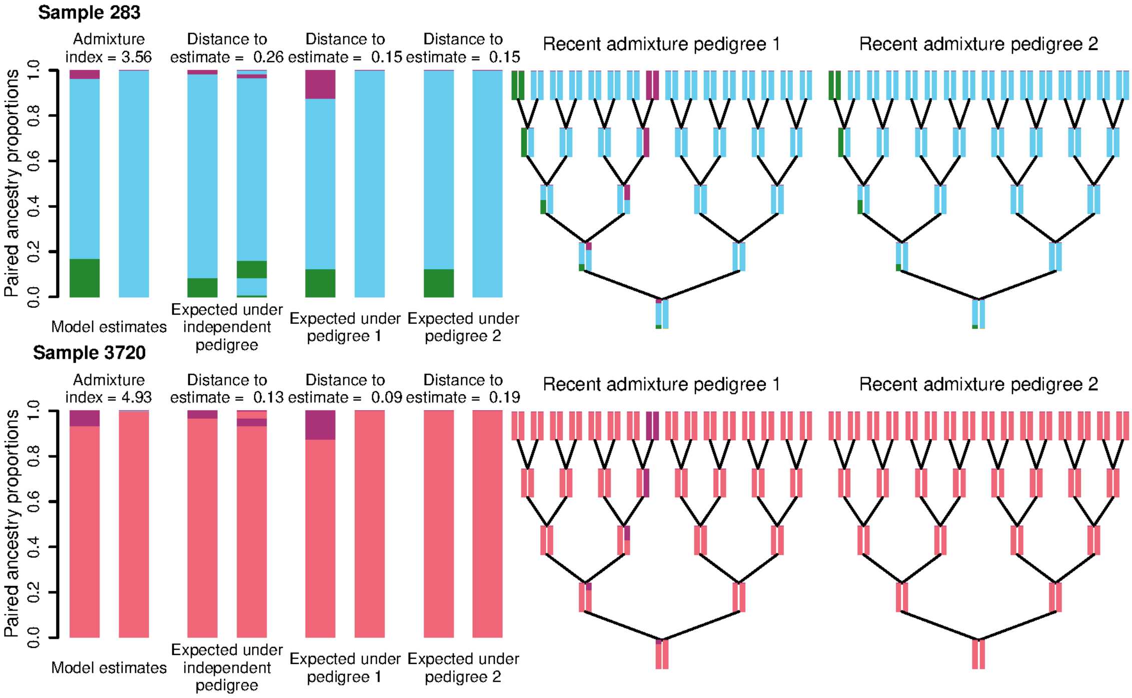


Figure S7: Ordered paired ancestry proportions and recent admixture pedigrees for the two Grant's gazelle samples with evidence of being recently admixed. Each row corresponds to a sample, where the left panel shows the ordered paired ancestries estimated with the parental admixture model, the ones expected under the independent pedigree and the ones expected under the two most compatible recent admixture pedigrees. Distances for the pedigrees are the Jensen-Shannon distance (JSD) to the model estimates. The right panel shows the two most compatible recent admixture pedigrees.
